# Lung CD4^+^ resident memory T cells use airway secretory cells to stimulate and regulate neutrophilic allergic airways disease

**DOI:** 10.1101/2024.10.03.616494

**Authors:** Vijay Raaj Ravi, Filiz T. Korkmaz, Carolina Lyon De Ana, Lu Lu, Feng-Zhi Shao, Christine V. Odom, Kimberly A. Barker, Aditya Ramanujan, Emma N. Niszczak, Wesley N. Goltry, Ian M.C. Martin, Catherine T. Ha, Lee J. Quinton, Matthew R. Jones, Alan Fine, Joshua D. Welch, Felicia Chen, Anna C. Belkina, Joseph P. Mizgerd, Anukul T. Shenoy

## Abstract

Neutrophilic asthma is a vexing disease, but mechanistic and therapeutic advancement will require better models of allergy-induced airway neutrophilia. Here, we find that periodic ovalbumin (OVA) inhalation in sensitized mice elicits allergic airway inflammation and pathophysiology mimicking neutrophilic asthma. OVA-experienced murine lungs harbor diverse clusters of CD4^+^ resident memory T_RM_ cells, including unconventional RORγt^negative/low^ T_H_17 cells. Acute OVA challenge instigates IL-17A secretion from these T_RM_ cells, driving CXCL5 production from Muc5ac^high^ airway secretory cells, leading to destructive airway neutrophilia. The T_RM_- and epithelial-cell signals discovered herein are also observed in adult human asthmatic airways. Epithelial antigen presentation regulates this biology by skewing T_RM_ cells towards T_H_2 and T_H_1 fates, so that the T_H_1-related IFN-γ suppresses IL-17A-driven, CXCL5-mediated airway neutrophilia. Concordantly, *in vivo* IFN-γ supplementation improves disease outcomes. Thus, using our model of neutrophilic asthma we identify lung epithelial-CD4^+^ T_RM_ cell crosstalk as a key rheostat of allergic airway neutrophilia.

**Highlights:**

- Recurrent OVA inhalation experience predisposes mice to allergic airways neutrophilia
- Neutrophil-prone lungs harbor CD4^+^ T_RM_ cells including RORγt^negative/low^ T_H_17 cells
- Muc5ac^high^ secretory cells instruct CD4^+^ T_RM_ fates and neutrophilia via MHC-II and CXCL5, respectively
- Prophylactic or therapeutic delivery of IFN-γ curbs allergic airway neutrophilia

## INTRODUCTION

Asthma is a chronic inflammatory disease of the respiratory tract characterized by goblet cell hyperplasia, mucus hypersecretion, and airway hyper-reactivity, which if unchecked can progress to irreversible airway plugging, tissue damage, and death ^1–3^. With ∼350 million people affected worldwide, asthma is a major health concern ^4^. Asthma can be eosinophilc or neutrophilic^5^. While eosinophilic asthma has been extensively modeled in research labs ^6,7^ leading to its recognition as a T helper 2 (T_H_2) cell driven, steroid-responsive pathophysiology ^1,3^, relatively less is understood about neutrophilic asthma which often presents as a more severe and steroid-resistant disease ^1,3,5,8^. Furthermore, some patients transition between phenotypes from treatment-responsive to treatment-resistant, for reasons yet unclear ^9–12^. These knowledge gaps underlie fewer therapeutic options and poorer quality of life for patients with neutrophilic asthma compared to their eosinophilic counterparts ^3^.

Natural and experimental exposures to inhaled allergens seed human and murine lungs with long-lived T_H_2- polarized CD4^+^ resident memory T (T_RM_) cells ^13–19^. These T_H_2 CD4^+^ T_RM_ cells localize around the airways, act as first line immune sentinels of the experienced lung ^15,19^, and produce more type-2 inflammatory cytokines like IL-5, and IL-13 compared to circulating counterparts. Consequently, T_H_2 T_RM_ cells induce hallmark features of eosinophilic asthma including eosinophil recruitment, mucus production, and airway hyper-reactivity ^15–19^. While these advances have elucidated host-factors underlying ‘T_H_2- high’ eosinophilic asthma and informed therapeutics targeting immune pathways for patients with this disease, key determinants of the more severe neutrophilic endotype of asthma remain uncertain.

A barrier to understanding neutrophilic asthma and the development of effective therapies is a lack of animal models ^3,20^. Traditional models of allergic asthma include sensitizing naïve mice to a prototypical allergen followed by pulmonary allergen challenge to induce T_RM_ cells and/or allergic inflammation ^6,7,15–19^. Such models robustly instigate T_H_2- high eosinophilic allergic airways disease, but are transient and weak inducers of neutrophilia. Exogenous inflammatory stimuli that elicit neutrophils and/or have T_H_17 adjuvanticity, like cyclic-di-AMP ^21^ or complete Freund’s adjuvant ^22–25^, have been added to the allergen exposures to make the allergic inflammation more neutrophilic. However, inciting neutrophils to an allergic response may not replicate allergy-induced neutrophilia, and such neutrophil-inducing adjuvants are unlikely to be triggers of human disease ^3,26^. Alternatively, since our airborne environment harbors endotoxins that may shape lung immunology ^27,28^, models combining LPS (either exogenously added or existing as contaminant in commercial reagents) with allergen exposures have been used to instigate allergic airways neutrophilia ^29–34^. These models have provided insights into IL-17A ^30^, IFN-γ ^21^, and G-CSF ^30^ in allergic airways, but translating findings from such models to patient settings has so far had limited success ^3,35,36^. Pertinent to this, although adult human lungs are known to be enriched for CD4^+^ T_RM_ cells ^14,37,38^, whether these lymphocytes have causative or mitigating roles in neutrophilic asthma is unclear. Adult humans have different lung CD4^+^ T_RM_ cells than children ^39^ and are disproportionately affected by neutrophilic asthma compared to children ^5,8^, with severity of neutrophilic inflammation predicting worse pulmonary outcomes ^40^. This suggests that the types of lung CD4^+^ T_RM_ cells may influence the asthma pathophysiology. CD4^+^ T_RM_ cells can influence lung epithelial cells to enhance neutrophilic inflammation during pneumonia ^41^, but if they do so in settings of allergy is uncertain. Furthermore, whether and how repeated environmental exposures to allergens remodels lung CD4^+^ T_RM_ cells and influences airway allergic disease is unclear.

Because recurrent but transient inhaled exposures to allergens are inherent to progression into adulthood, we hypothesized that multiple intermittent bouts of inhaled allergens would induce progression of allergic airways disease from eosinophilic to neutrophilic. With that rationale, we establish a mouse model of neutrophilic asthma that resulted from repeated allergen exposures, involving the physiologically relevant route of recurrent exposure (inhalation), inbred animals for which the greatest variety of genetically engineered lines are available (C57BL/6 mice), and a simple model allergen for which a large panoply of antigen-specific immunological tools have been developed (ovalbumin). Thus, our approach elicits allergy-induced airway neutrophilia and its resultant pathophysiology within a useful, tractable, and forward-driving animal model. This model allowed us to comprehensively phenotype CD4^+^ T_RM_ cells in the experienced allergic lungs, to highlight an unconventional subset of pathogenic T_H_17 T_RM_ cells, and to use genetically engineered mice to elucidate an epithelium-lymphocyte-neutrophil communication triad underlying allergic airways neutrophilia. By identifying key cellular and molecular contributors and constrainers of neutrophilic allergic airways disease, we were guided to identify IFN-γ as a potent suppressor of neutrophilic allergic airways disease. Our results now support further investigation of recombinant IFN-γ and/or the pathways this cytokine triggers as approaches to countering neutrophil asthma.

## RESULTS

### Transient and recurrent allergen experience begets allergic airway neutrophilia

Relevant environmental exposures make laboratory mice better resemble adult humans in regard to immune cell localizations and activities ^42–47^. Conventional ovalbumin (OVA)-induced animal model of allergic asthma, relying on sensitized mice being acutely challenged with aerosolized OVA, capture phenotypes of T_H_2 asthma (**Fig S1A**) including eosinophilic influx with negligible neutrophilia (**Fig S1B,** Gating Strategy in **Fig S2**). Since this model does not account for aeroallergen experiences which establish and instruct the CD4^+^ T_RM_ cells around human airways ^5,8,14^, we hypothesized that repeated and intermittent exposures of sensitized mice to inhaled allergens over extended duration will better reflect the human experience ^48^ and yield allergic airways disease more characteristic of the neutrophilic asthma so currently challenging. C57BL/6J mice were sensitized to OVA in the typical fashion but then administered intermittent recurrent exposures to aerosolized OVA allergen (or PBS vehicle as a negative control) separated by multiple weeks of recovery (to avoid induction of tolerance^49^), before a final challenge with OVA to induce disease exacerbation across all the sensitized mice (**Fig 1A**). Sets of mice with identical sensitizations and final challenges but differing in inhaled exposures to allergen will hereafter be referred to as Lung History (LH) or No Lung History (No-LH) groups. Intranasal challenge with OVA induced more severe disease in the LH mice when compared to No-LH mice. LH mice showed severe sickness behavior within 4 hours of OVA challenge (**Fig S1E**). They also exhibited rapid and severe (>5%) loss in body weight within 24 hours; this loss persisted through at least 48 hours (**Fig S1F,G**). Due to the severity of the disease elicited in LH mice, we hereafter picked the 24 hour (or earlier) timepoints for our challenge studies unless otherwise stated.

**Figure 1:**
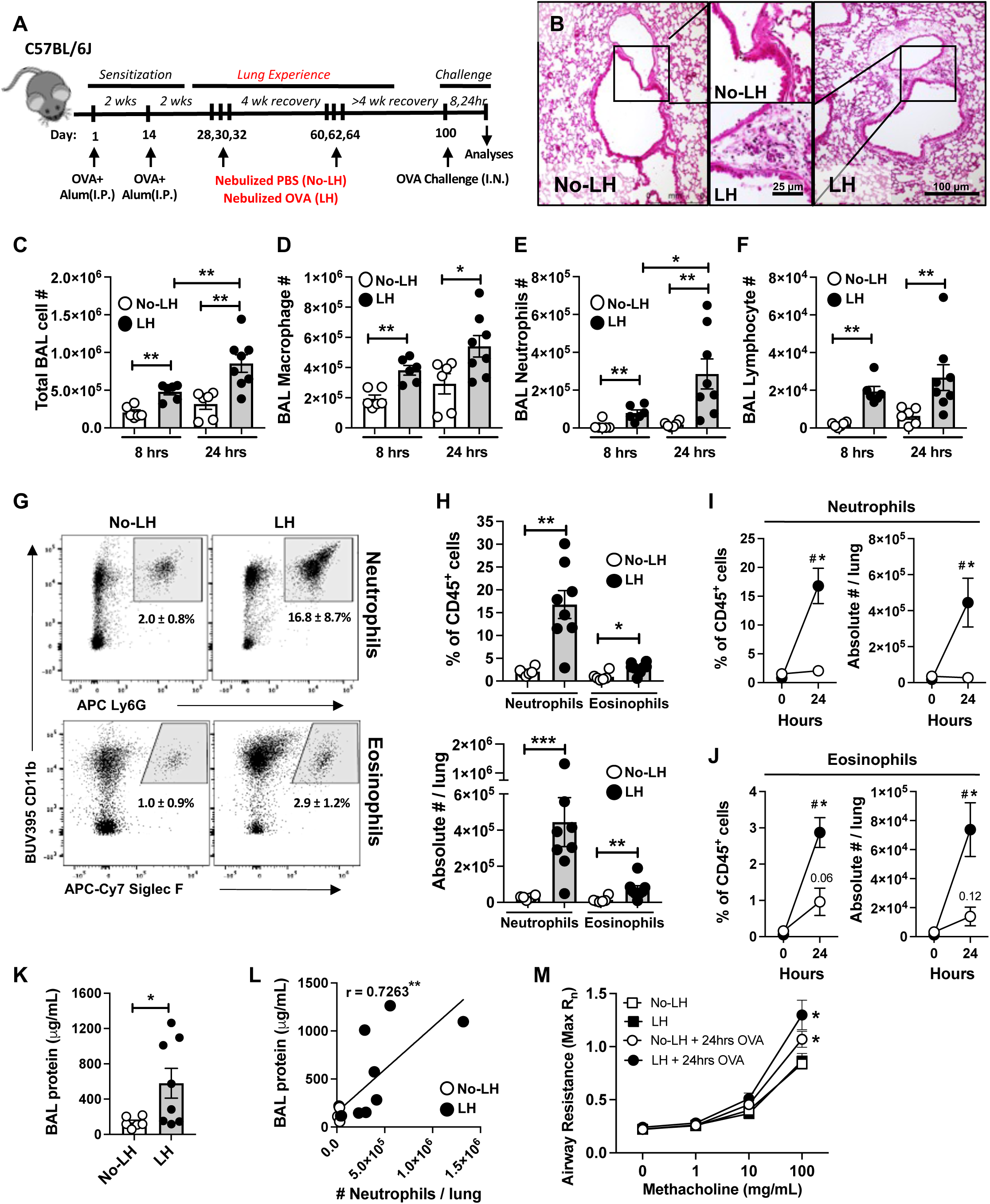
Mouse model of neutrophilic asthma. **A)** Schematic of experimental timeline used. **B)** Representative images of hematoxylin and eosin stained sections from mice (n=3 mice) 8 hours post OVA challenge. **C)** Total cell-, **D)** macrophage- **E)** neutrophil-, and **F)** lymphocyte-numbers in bronchoalveolar lavages (BAL) from mice at designated timepoints post OVA challenge. Mann-Whitney test. **G)** Representative dot plots depicting lung (ivCD45.2^-^) neutrophils and eosinophils as fraction (%) of CD45^+^ cells 24 hours post OVA challenge, mean ± SD. **H)** Numbers of lung (ivCD45.2^-^) neutrophils and eosinophils 24 hours post OVA challenge. Mann-Whitney test. **I)** Numbers of lung (ivCD45.2^-^) neutrophils and **J)** eosinophils in no-LH (white dots) and LH mice (black dots) at baseline and 24 hours post OVA challenge. Two-way ANOVA with Fisher’s LSD Test. **K)** Lung damage expressed as BAL protein content. Mann-Whitney test. **L)** Scatterplot correlating lung neutrophil numbers and lung damage 24 hours post OVA challenge. Spearman’s correlation coefficient (r) and statistical significance denoted. **M)** Airway reactivity in response to increasing doses of methacholine in mice at baseline and 24 hours post OVA challenge as measured by Flexivent. The measurements represent Maximum Rn values. Two-way ANOVA with Fisher’s LSD Test. All data have n≥5 mice, 2-3 experiments, mean ± SEM, *p* value: *≤ 0.05, **≤ 0.01, ***≤ 0.001.

Next, we sought to characterize the inflammatory profiles of the mouse airways. Consistent with rapid onset of severe disease, LH mice displayed more pronounced peribronchial inflammation (**Fig 1B**) characterized by elevated bronchoalveolar lavage (BAL) cellularity (**Fig 1C**) that included greater macrophage (**Fig 1D**), neutrophil (**Fig 1E**), and lymphocyte (**Fig 1F**) numbers within 8 hours of challenge when compared to No-LH mice. Flow cytometry using intravital staining to exclude circulating leukocytes also revealed greater frequencies and numbers of extravascular (ivCD45.2^-^) lung neutrophils (**Fig 1G,H**), with a modest contribution of eosinophils (**Fig 1G,H**) in the LH mice compared to their No-LH counterparts. Of note, prior to allergen challenge, both LH and No-LH mice had negligible and comparable numbers of extravascular neutrophils and eosinophils in the lung (**Fig 1I,J**), indicating complete resolution of prior inflammation. Furthermore, while allergen exposure increased both neutrophils and eosinophils in the LH mice (**Fig 1I**), the lungs of No-LH mice displayed modest numerical increases that were confined to eosinophils (**Fig 1J**).

We considered several limitations. Endotoxin is a human-relevant environmental exposure of interest ^27,28^ and independently capable of triggering neutrophil recruitment. So, the robust airway neutrophilia observed in LH mice could result from endotoxin contamination within our OVA preparations ^22^. Such concerns are partially alleviated by the fact that both LH and No-LH mice were challenged with the same OVA preparations but exhibited different outcomes based on lung history (**Fig 1A-J**). Furthermore, naïve mice with no prior OVA sensitization showed negligible neutrophilic infiltration after OVA challenge (**Fig S1L**), suggesting any possible endotoxin contamination of the OVA preparation was insufficient to induce the robust neutrophilia observed in the LH mice. Another concern is timing. In the standard murine models of OVA-specific allergic airways, a modest and transient neutrophilic infiltration occurs, but eosinophils then become predominant after the allergen challenge^50,51^. In our study, mice without inhalation histories reflected such cellular dynamics (**Fig S1I-J**). In contrast, our LH mouse model was distinct in that both neutrophils and eosinophils were elevated compared to No-LH controls at all time-points, and the airway neutrophilia in the LH mice was consistently dominant throughout the time-course, persisting through at least 48 hours post OVA challenge (**Fig S1I-J**). Thus, the inflammatory response including robust airway neutrophilia seen in LH mice differs dramatically from that observed in mice without inhalation histories, reflective of immunological remodeling induced by inhaled allergen experience. Indeed, OVA challenge of sensitized mice ushered through the conventional model of eosinophilic asthma (i.e., without lung history) displayed very significantly robust eosinophilic, but weaker neutrophilic, influxes within 24 hours when compared to PBS challenge (**Fig S1M**), an eosinophilic response comparable to that observed in LH mice (**Fig 1H** and **Fig S1M**).

Increased vascular permeability and lung edema with plasma leakage into the airways is a feature of asthma ^52^. Consistent with the detrimental nature of neutrophilic inflammation, LH mice demonstrated greater airway edema (measured as BAL protein content, **Fig 1K**) which positively correlated with airway neutrophilia at 24 hours (**Fig 1L**), and remained high at 48 hours (**Fig S1K**). Another key clinical feature we found was reproducible in our mouse model of neutrophilic asthma was its steroid-resistance (**Fig S3**) ^1,3,5^. We found that the allergic airways inflammation (**Fig S3A-C**), mechanisms driving neutrophilic influx (**Fig S3D**, to be discussed further) and the resulting airway damage (**Fig S3E**) remained refractory even to high dose prophylactic dexamethasone therapy in LH mice. Finally, both LH and No-LH mice exhibited comparable airway resistance at baseline and presented with exacerbated airway hyper-reactivity to methacholine after inhaled allergen memory recall challenge (**Fig 1M**). This observation coupled with its capacity for rapid eosinophilia (**Fig 1J**) confirms the utility of sensitized mice without inhaled allergen history (i.e. No-LH mice) as a model of eosinophilic asthma. More importantly, our results demonstrate that transient and recurrent exposure of sensitized mice to inhaled ovalbumin mimics key features of human neutrophilic asthma, yielding a useful model of this morbid disease.

### Allergic airway neutrophilia is accompanied by multiple myeloid cell changes

Traditional models of eosinophilic asthma as in **Figure S1A** are accompanied by changes to the myeloid landscape involving alveolar macrophages ^53^, monocytes ^54^, CD11b^+^ dendritic cells (DCs)^55^, and eosinophils ^56^. To determine whether myeloid cells were different beyond the neutrophil subset in the LH model evoking neutrophilia, we used flow cytometry analyses of single cell suspensions from lungs of mice in whom the intravascular leukocytes were discriminated via *in vivo* staining from intravenous injection of anti-CD45.2 antibody (**Fig S4,** Gating Strategy in **Fig S2**). At baseline (prior to acute allergen challenge), the LH mice had elevated numbers of tissue resident interstitial macrophages **(Fig S4B)**, but other lung myeloid cell numbers were comparable to the No-LH mice. Allergen challenge induced elevated recruitment of monocytic cells in LH mice observed as accumulation of monocyte-derived interstitial macrophages (**Fig S4D**), Ly6C^+^ inflammatory monocytes (**Fig S4E**), and Ly6C^-^ patrolling monocytes (**Fig S4F**) while instigating a modest decrease in numbers of monocyte-derived alveolar macrophages (**Fig S4C**). Among the DCs, CD11b^+^ cDCs have pathogenic functions in eosinophilic asthma ^57,58^, while plasmacytoid DCs play protective roles ^59–61^. Both these cell types were expanded in LH mice (**Fig S4G,H**) while no changes in CD103^+^ cDCs (**Figs S4I**) or CD103^+^CD11b^+^ cDCs (**Figs S4J**) were observed in the LH lung after allergen challenge. These results together reveal that allergen-experienced lungs have multiple diverse alterations in lung myeloid cells alongside the distinct accumulation of neutrophils.

### Lung cell-derived CXCL5 associates with allergic airway neutrophilia

Considering the neutrophil attracting chemokines within the lungs of LH and No-LH mice after allergen challenge, both groups showed comparable CXCL1 (**Fig S5A**) and CXCL2 (**Fig S5B**) but the LH airways were enriched for CXCL5 (**Fig S5C**) and diminished for CXCL10 (**Fig S5D**). CXCL5 correlated with lung neutrophil numbers and BAL protein, but the other CXC chemokines did not (**Fig S5E-L**). To determine the cellular sources of chemokines and how the distinct cells sources were impacted by prior lung allergen history, we measured chemokine transcripts in sorted CD45^+^ leukocytes, CD45^-^EpCAM^+^ epithelial cells, and CD45^-^EpCAM^-^ stromal cells (which includes mesenchymal and endothelial cells) from allergen-challenged lungs of LH and No-LH mice. CXCL5 was more strongly expressed in lung epithelial and stromal cells (**Fig S5O**), consistent with other settings of neutrophilic pulmonary inflammation ^41,62^. Compared to No-LH mice, the LH mice had higher CXCL5 message in both lung epithelial cells and lung stromal cells, but not lung leukocytes (**Fig S5O**). In contrast, none of the other chemokines were increased in any cell-type due to prior inhaled allergen experiences (**Fig S5M,N,P**). Thus, elevated CXCL5 expression by lung epithelial and stromal cells uniquely associates with the airway neutrophilia in allergen experienced mice.

### Lungs of mice with inhaled allergen history harbor diverse CD4^+^ T_RM_ cells

Respiratory exposure to inhaled antigens lead to the formation of regionally compartmentalized lung-resident CD4^+^ memory T_RM_ cells that on subsequent memory recall secrete lineage specific cytokines to orchestrate rapid innate immunity ^37^. T_H_17 T_RM_ cell-derived IL-17A augments epithelial CXCL5 production to accelerate neutrophilic inflammation in the lungs during pneumonia ^41,63,64^. Given the rapidity of neutrophil responses (**Fig 1E**), we examined IL-17A and CXCL5 at the early time-point of 8 hours after allergen challenge. Both cytokines were already elevated (**Fig 2A-B**), as was BAL protein characteristic of lung edema **(Fig 2C)** in the LH compared to No-LH mice. Strong positive correlations were observed among these inflammatory outcomes **(Fig 2D).** We considered whether T_H_17 T_RM_ cells might also be present in the lungs of mice predisposed to neutrophilic allergic airways disease. Consistent with CD4^+^ T_RM_ cell deposition due to allergen experience, lungs of LH mice were enriched for extravascular CD11a^high^CD69^+^ CD4^+^ T cells (**Fig 2E,F**, Gating Strategy in **Fig S6**) that were negative for CD62L and high for CD44 expression (**Fig 2G**). In contrast, mice ushered though the traditional model of eosinophilic asthma exhibited activated CD4^+^ T cells in their lungs during eosinophilic inflammation but no CD4^+^ T_RM_ cells at baseline (**Fig S1C,D**). Thus, CD4^+^ T_RM_ cells are a salient feature of lungs predisposed to neutrophilic allergic airways disease.

**Figure 2:**
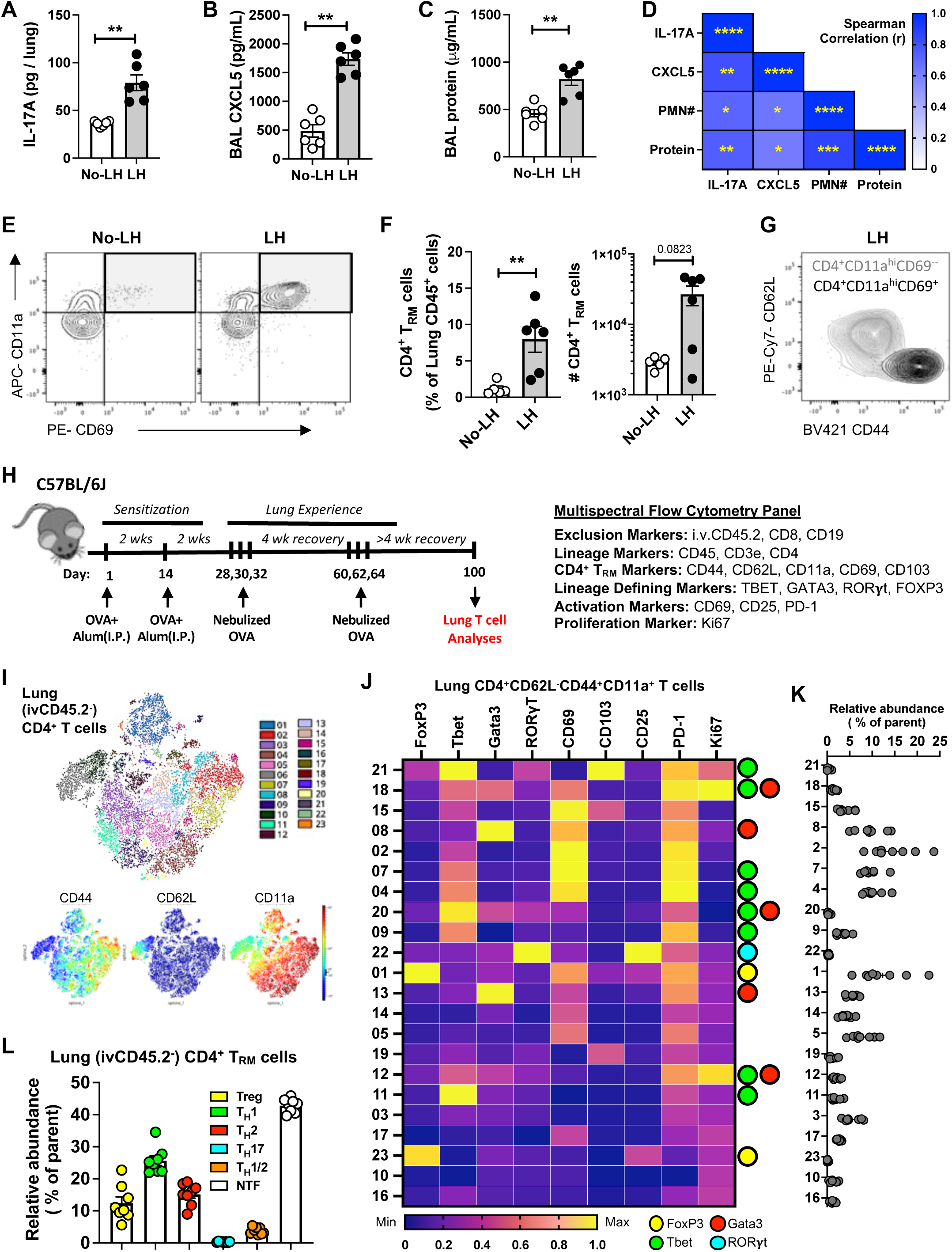
Diverse clusters of tissue resident CD4^+^ T_RM_ cells reside in lungs of mice with inhaled allergen history. Levels of **A)** whole lung IL-17A, **B)** BAL CXCL5 and **C)** lung damage in mice 8 hours post OVA challenge. Mann-Whitney test. **D)** Heat map depicting Spearman’s correlation coefficients (r) and statistical significance for the specified correlations. **E)** Representative contour plots depicting lung (ivCD45.2^-^) CD4^+^ T_RM_ cells identified as CD69^+^CD11a^high^ CD4^+^ T cells in unchallenged mice. **F)** Numbers of lung (ivCD45.2^-^) CD4^+^ T_RM_ cells in unchallenged mice. Mann-Whitney test. **G)** Representative contour plots illustrating CD62L and CD44 levels on lung (ivCD45.2^-^) CD4^+^ T_RM_ cells (identified as CD69^+^CD11a^high^ CD4^+^ T cells) in comparison to CD69^-^CD11a^high^ CD4^+^ T cells in unchallenged LH mice. **H)** Schematic of experimental timeline and antibody panel used. **I)** Phenograph clustering overlaid on opt-SNE projection depicting lung (i.v.CD45.2^-^) CD4^+^ T cells concatenated from n=8 LH mouse lungs on day 100. opt-SNE projection with heatmap visualization depicting CD44, CD62L and CD11a expression levels are shown. **J)** Heat map depicting normalized expression levels of distinct molecules on lung (ivCD45.2^-^) effector memory (CD62L^-^CD44^+^CD11a^+^) CD4^+^ T cell clusters. Lineage determining transcription factor (LDTF) status of each cluster is depicted on the *right*. **K)** Relative abundance of each cluster in LH mice at day 100 is shown. **L)** Frequencies of distinct memory T_H_ cell lineages within lungs of LH mice at day 100. Positivity for a tested LDTF was determined using cutoffs identified from clusters negative for that LDTF. *p* value: *≤ 0.05, **≤ 0.01, ***≤ 0.001, ****≤ 0.0001. All data have n≥5 mice, 2 experiments, mean ± SEM.

We phenotyped the CD4^+^ T_RM_ cells in experienced lungs to identify helper T cell subsets discriminated by lineage defining transcription factors (LDTFs), including RORγt which denotes and mediates T_H_17 cell biology ^65^. We designed a 21-parameter high-dimensional multispectral flow cytometry (MSFC) panel which probed for LDTFs as well as markers for CD4^+^ T_RM_ cell subsets, activation, and proliferation (**Fig 2H**). To achieve unbiased identification and quantification of CD4^+^ T_RM_ cell subsets, we computationally concatenated extravascular (ivCD45.2^-^) CD4^+^ T cells from lungs of 8 LH mice, projected the data into 2-dimensional space using optimized t-distributed stochastic neighbor embedding (opt-SNE) ^66^, and clustered the data in unsupervised fashion using the Phenograph algorithm ^67,68^. At least 23 distinct clusters of CD4^+^ T cells were observed (numbered in order of decreasing abundance), highlighting an unforeseen complexity within the CD4^+^ T cell pool of lungs susceptible to allergic airway neutrophilia (**Fig 2I**). Among these, cluster 6 was naïve CD4^+^ T cells based on CD62L^+^CD44^-^ staining (**Fig 2I**), so excluded from further analyses to focus on lung memory ivCD45.2^-^CD62L^-^CD44^+^CD11a^+^CD4^+^ cells **(Fig 2J)**. We found two FoxP3^+^ clusters (1 and 23), five Tbet^+^ clusters (4, 7, 9, 11, and 21), two Gata3^+^ clusters (clusters 08 and 13), and three Tbet^+^Gata3^+^ clusters (12,18, and 20)**(Fig 2J,K)**, consistent with distinct Treg, T_H_1, T_H_2, and polyfunctional T_H_1/T_H_2 phenotypes, respectively. Only one cluster (cluster 22) was RORγt^+^ and consistent with conventional T_H_17 cells. Many lung CD4^+^ T cells in LH mice lacked all four LDTFs tested (9 of the 22 clusters: 2, 3, 5, 10, 14, 15, 16, 17, and 19) **(Fig 2J,K).** This was surprising, since IL-17A led us to anticipate T_H_17 T_RM_ cells. Cumulative frequencies revealed that T_H_1, T_H_2, Treg and LDTF-negative T_RM_ cells were common, but T_H_17 T_RM_ cells were vanishingly low **(Fig 2L)**. This contrasts with our studies using the same approach and lungs recovered from pneumococcal pneumonia, in which RORγt^+^ T_H_17 T_RM_ cells are abundant ^69^. As in prior studies ^69^, the blood (ivCD45.2^+^) CD4^+^ T cell subset distributions and surface marker phenotypes differed from the lungs; the blood also lacked RORγt^+^ cells in the LH mice **(Fig S7A-D)**. While the No-LH mice failed to accumulate significant lung CD4^+^ T_RM_ cells (**Fig 2E,F**), they possessed circulating CD4^+^ T cells (**Fig S7E**), CD4^+^ T_CM_ cells (**Fig S7F**), and CD4^+^ T_EM_ cells (**Fig S7G**) with similar phenotypes (**Fig S7D**) compared to LH mice, which reflects the shared experience of systemic sensitization. Thus, although acute disease exacerbation involved robust IL-17A production in the lungs of LH mice, conventional RORγt^+^ T_H_17 T_RM_ cells were not present in these lungs.

### Unconventional T_H_17 T_RM_ cells are sources of IL-17A that drive allergic airway neutrophilia

Because the rapid appearance of IL-17A in lungs without RORγt^+^ CD4^+^ T_RM_ cells was surprising, we sought to determine whether and which lymphocytes from these lungs might be poised to produce IL-17A. We stimulated single cell suspensions from lungs of LH and No-LH mice *ex vivo,* which induced IL-17A expression from both CD4^-^ and CD4^+^ T cells (**Fig 3A,B**). While CD4^-^ T cells were the predominant source of IL-17A in the No-LH lungs (**Fig 3B**), the majority of the IL-17A producers in the LH lungs were CD4^+^ T cells (**Fig 3B**). This was supported by significant enlargement in IL-17A-producing CD4^+^ T cell pool in the LH lungs (**Fig 3C**). Surprisingly, these IL-17A^+^ CD4^+^ T cells from LH lungs expressed less RORγt than the CD4^-^ T cells from the same lungs or the CD4^+^ T cells from lungs of mice recovered from *S. pneumoniae* infections (**Fig 3D**)^69^ despite comparable IL-17A expression (**Fig 3E**). Furthermore, these IL-17A-producing CD4^+^ T cells from LH lungs were distinct from the other traditional helper cell subsets, in that they did not contain Tbet or Gata3 nor did they co-express IFN-γ, IL-5, or IL-13 (**Fig S8A,B**). While the blood of LH mice also possessed CD4^-^ and RORγt^negative/low^ CD4^+^ T cells that could produce IL-17A (**Fig S8C,D**), their numbers were comparable to blood of No-LH mice (**Fig S8E**) which responded with poor IL-17A secretion on memory recall challenge *in vivo* (**Fig 2A**). These findings suggest that lungs (but not blood) of mice with allergen experience are enriched for RORγt^negative/low^ T_H_17 T_RM_ cells that promptly secrete IL-17A to drive CXCL5 secretion and allergic airways neutrophilia upon activation. Indeed, depletion of CD4^+^ cells just before final allergen challenge (**Fig S9A**) or genetic ablation of IL-17A/F (**Fig 3F**) compromised CXCL5 accumulation (**Fig 3G, Fig S9B**) and allergic neutrophilia **(Fig 3H, Fig S9C,D)**, but not eosinophilia (**Fig 3I**), in the airways of LH mice despite their extensive inhalation experience. Furthermore, consistent with a role of CD4^+^ T cell driven antigen specificity and not just generalized innate immune memory in this biology, challenge of mice with an irrelevant antigen did not exacerbate airway neutrophilia (**Fig S10**).

**Figure 3:**
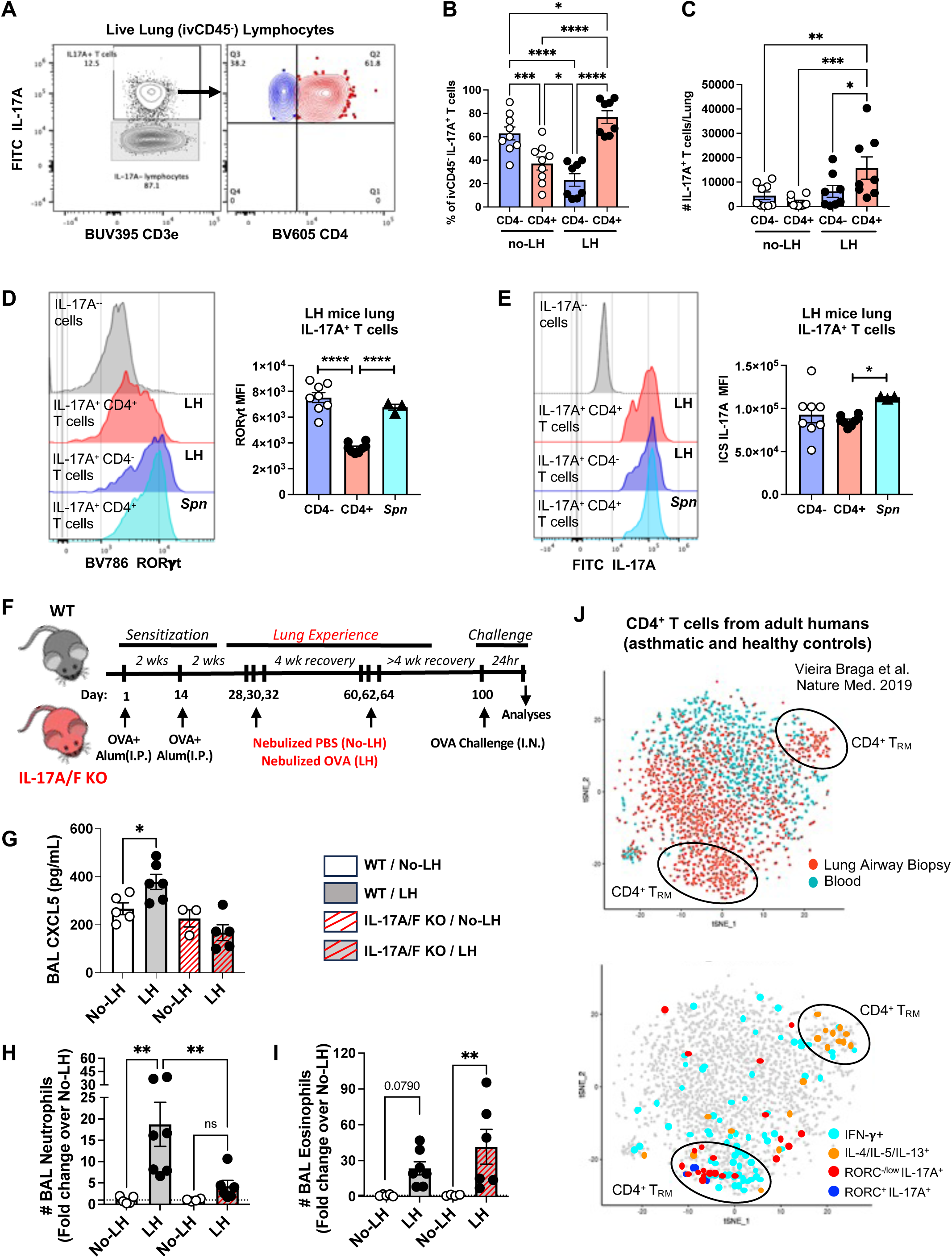
Allergen-experienced lungs include unconventional RORγt^negative/low^ T_H_17 T_RM_ cells that drive allergic airway neutrophilia. **A)** Representative contour plots depicting IL-17A-producing CD4^-^ and CD4^+^ T cells as identified from live lung (ivCD45.2^-^) lymphocytes within single cell suspensions of LH mouse lungs stimulated with PMA/ionomycin. **B)** Relative frequency of IL-17A^+^ lung (ivCD45.2^-^) CD4^-^ and CD4^+^ T cells in No-LH and LH mice. One-way ANOVA with Fisher’s LSD Test. **C)** Absolute numbers of IL-17A^+^ lung (ivCD45.2^-^) CD4^-^ and CD4^+^ T cells in No-LH and LH mice. One-way ANOVA with Fisher’s LSD Test. **D)** Representative histograms and expression levels of RORγt in IL-17A producing lung (ivCD45.2^-^) CD4^-^ and CD4^+^ T cells in LH mice. One-way ANOVA with Fisher’s LSD Test. **E)** Representative histograms and expression levels of IL-17A in IL-17A producing lung (ivCD45.2^-^) CD4^-^ and CD4^+^ T cells in LH mice. One-way ANOVA with Fisher’s LSD Test. Expression patterns of *S. pneumoniae* specific lung (ivCD45.2^-^) CD4^+^ T cells included as positive control for conventional T_H_17 signatures. **F)** Schematic of experimental timeline used. **G)** Bronchoalveolar lavages (BAL) CXCL5 in WT and IL-17A/F knockout No-LH and LH mice 24 hours post OVA challenge. One-way ANOVA with Fisher’s LSD Test. Total **H)** neutrophil and **I)** eosinophil numbers in BALs from WT and IL-17A/F knockout No-LH and LH mice 24 hours post OVA challenge. One-way ANOVA with Fisher’s LSD Test. *p* value: *≤ 0.05, **≤ 0.01, ***≤ 0.001, ****≤ 0.0001. All data have n≥5 mice, 2 experiments, mean ± SEM. **H)** t-SNE projection of scRNA-Seq data depicting expression patterns for T_H_ cell signature markers of interest as expressed by CD4^+^ T cell isolated from airway wall biopsies and peripheral blood of adult asthmatic and healthy humans as displayed on the interactive web portal: https://asthma.cellgeni.sanger.ac.uk/ ^14^.

Given these findings, we sought to mine independent single cell RNA-sequencing (scRNA-Seq) databases from asthmatic human-^14,70^ and allergic murine-airways^17^ for presence of the unconventional RORγt^negative/low^ T_H_17 T_RM_ cells. Consistent with our findings in LH mice, interrogation of a scRNA-Seq dataset profiling CD4^+^ T cells from airway wall biopsies and peripheral blood samples of adult asthmatic and healthy humans^14^ showed that the airways (and not blood) of adult humans are also enriched for RORγt^negative/low^ T_H_17 T_RM_ cells in addition to T_H_1, T_H_2 and Treg T_RM_ cells (**Fig 3J, Fig S11**) and neutrophils but not eosinophils ^14^. Reanalyses of a separate scRNA-Seq dataset of adult asthmatic and healthy human airways 24 hours post segmental allergen challenge^70^ also confirmed the presence of RORγt^negative/low^ T_H_17 T_RM_ cells in human airways (**Fig S12A-C**). Of note, the unconventional T_H_17 T_RM_ cells that were enriched in our mouse model (∼6% of all lung CD4^+^ T cells) and human airways (>2% of all lung CD4^+^ T cells, **Fig S12C**) were vanishingly scant in murine airways with allergic airway eosinophilia^17^ (<0.8% of airway CD4^+^ T cells, **Fig S12D-F**). Thus, adult human lungs contain T_H_17 T_RM_ cells prone to IL-17A expression despite little to no RORγt, as were identified in mice with a history of inhaled allergen exposures and a predisposition to neutrophilic allergic airways disease.

### Airway Muc5ac^high^ secretory epithelial cells communicate with CD4^+^ T cells and neutrophils

Lung epithelial cells use MHC-II to function as antigen presenting cells (APCs) for CD4^+^ T cells during pneumonia ^69^. Roles and dynamics of epithelial MHC-II in allergic lung disease are unknown. We examined professional APC-related molecules on epithelial cells isolated from LH mice at baseline and 24 hours after allergen challenge. Mice expressing GFP under control of human surfactant protein C promoter helped distinguish cell-types (**Fig S13A**) ^71^ including SPC^low^MHC^high^ alveolar epithelial cells that are the highest MHC-II expressors in resting mouse lungs ^69^. At baseline, alveolar epithelial cells had higher expression of MHC-II, as expected ^69^, while epithelial cells from the conducting airways tended to have higher expression of the other molecules examined (**Fig 4A, Fig S13B**). Allergen challenge of LH lungs enhanced expression of all the tested molecules in secretory cells from the airways **(Fig 4B, Fig S13B)**, which includes club cells and goblet cells. Type 2 alveolar epithelial cells increased expression of MHC-II, CD80, and CD86 after allergen challenge, whereas no changes were observed for multiciliated cells or SPC^low^MHC^high^ cells after the allergen challenge **(Fig 4B, Fig S13B)**. The airway secretory cells were the most consistently changed epithelial subset due to allergen challenge (**Fig 4B**), which is of interest since asthma pathophysiology particularly involves conducting airways and their secretory cells. In the allergen-challenged airways, secretory cells were the sole expressers of Muc5ac (**Fig 4C**), as expected. In addition, these cells were the only epithelial source of CXCL5 (**Fig 4D**), the neutrophil-attracting chemokine that was exacerbated by inhaled allergen history (**Fig 2B, Fig S5**). These findings suggest that Muc5ac^high^ secretory epithelial cells organize immunopathologic niches containing CD4^+^ T_RM_ cells and neutrophils around allergic airways during neutrophilic asthma. Indeed, immunofluorescence analyses of LH lungs 8 hours post allergen challenge revealed biased localization of CD4^+^ cells and neutrophils near the airway, but not the alveolar, epithelium (**Fig 4E**). Furthermore, exploration of scRNA-seq data profiling airway epithelial cells from adult asthmatic and healthy human lungs confirmed murine findings and revealed that Muc5ac^high^ airway secretory cells from human subjects (**Fig 4F**) also expressed MHC-II (**Fig 4G, Fig S13C)**, MHC-II related accessory and costimulatory molecules (**Fig 4H, Fig S13D,E)**, IL-17A receptor components (**Fig S13F)**, and CXCL6 which is the human ortholog of murine CXCL5 (**Fig 4I**) ^14,72^. Muc5ac^high^ secretory cells in adult human airways expressed very little to no inhibitory accessory molecule HLA-DO (**Fig S13D)** suggesting active antigen presentation in these cells at time of isolation. Also, CD80, CD86 and PD-L2 expression was minimal suggesting species-specific differences in airway epithelial biology (**Fig S13E).** Of note, pan-epithelial survey of scRNA-seq data from the same study confirmed that Muc5ac^high^ airway secretory cells were high expressors of these immune facing molecules among all the epithelial cells within adult human lungs (**Fig S14**) ^14^ and cross-comparison of these signals in a different adult human lung scRNA-Seq dataset also confirmed our findings (**Fig S15**)^73^. Thus, as the only epithelial cells showing elevations across the antigen presentation proteins measured, and the primary epithelial source of the neutrophil chemokines CXCL5/CXCL6, the Muc5ac^high^ airway secretory cells bridge CD4^+^ T_RM_ cells and neutrophils in lungs with extensive inhalation histories.

**Figure 4:**
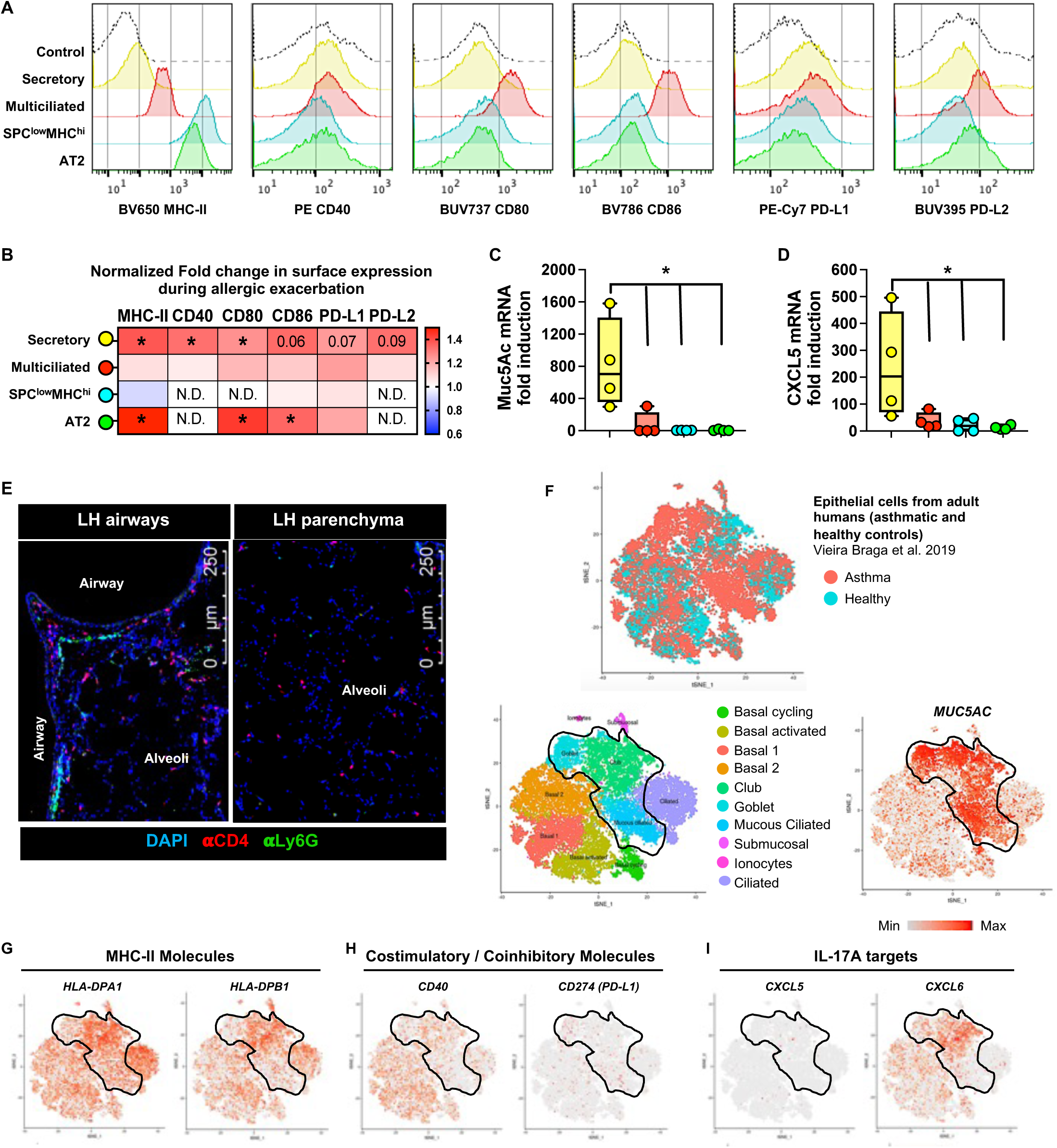
Muc5ac^high^ airway secretory cells communicate with CD4^+^ T cells and neutrophils in inhaled allergen experienced lungs. **A)** Representative histogram plots depicting surface expression patterns of MHC-II, costimulatory molecules CD40, CD80, CD86 and coinhibitory molecules PD-L1 and PD-L2 on distinct epithelial cells from LH mice at baseline. **B)** Heat Map depicting fold change in surface expression levels of specified APC-related molecules on distinct epithelial cells from LH mice 24 hours post OVA challenge normalized to their baseline counterparts. N.D., not detected above FMO control. Mann-Whitney test. **C, D)** mRNA levels of **C)** Muc5ac and **D)** CXCL5 in FACS-sorted epithelial cells isolated from LH mice 24 hours post OVA challenge. One-way ANOVA with Fisher’s LSD Test. *p* value: *≤ 0.05. All data have n=4 mice, 2 experiments, mean ± SEM. **E)** Representative immunofluorescent micrographs showing anatomical location of Scgb1a1^+^ secretory epithelial cells (green), CD4^+^ T cells (*red*), and Ly6G^+^ neutrophils (magenta) in LH lungs 8 hours post OVA challenge. DAPI (blue) used as a counterstain to identify lung structures. Data represents *n* = 3 mice/timepoint, two experiments. **F-K)** t-SNE projection of scRNA-Seq data depicting expression patterns for designated markers of interest by different subsets of epithelial cells identified in airways of adult asthmatic and healthy humans as displayed on the interactive web portal: https://asthma.cellgeni.sanger.ac.uk/ ^14^.

### Antigen presentation by epithelial cells governs CD4^+^ T_RM_ cell activities

To test whether epithelial antigen presentation impacts CD4^+^ T_RM_ cells in the lungs with inhaled allergen history, we studied LH mice lacking MHC-II specifically in lung epithelial cells ^69^, referred to as MHC-II^ΔEpi^ mice after tamoxifen-induced gene targeting. MHC-II^ΔEpi^ mice and similarly tamoxifen-treated but Cre-negative littermate MHC-II^fl/fl^ mice were ushered through the regimen of sensitization and inhaled allergen exposures, after which lung CD4^+^ T_RM_ cells were phenotyped **(Fig 5A)**. No differences in abundance of CD4^+^ T_RM_ cells were observed between genotypes **(Fig S16A)**. Phenograph clustering of concatenated lung CD4^+^ T_RM_ cells (ivCD45.2^-^CD62L^-^CD44^+^CD11a^+^) identified 20 distinct clusters **(Fig 5B,C),** none of which were unique to either genotype. Only 1 small RORγt^+^ cluster (cluster 19) was observed **(Fig 5C-D)**, which in these lungs also expressed Gata3^+^ (T_H_2/17). Deletion of epithelial MHC-II very modestly perturbed CD4^+^ T_RM_ cell abundances on a per cluster level (**Fig 5D and Fig S16B**). However, enumeration of CD4^+^ T_RM_ cells sharing common helper T cell phenotypes (based on expression of individual LDTFs) revealed significant reductions in Tbet^+^ CD4^+^ T_RM_ cells, except for those coexpressing Foxp3, which instead increased **(Fig 5E)**. Because Tbet^+^ Foxp3^+^ Treg cells potently suppress Tbet-dependent T_H_1 responses ^74^, our results suggest that deletion of epithelial MHC-II leads to an expansion of Treg T_RM_ cells to suppress T_H_1 T_RM_ cell activity. Indeed, correlation analyses revealed strong inverse correlation between these CD4^+^ T cell types (**Fig S16C**) and *ex vivo* stimulation revealed 30% reduction in IFN-γ secreting lung CD4^+^ T cells (**Fig 5F**). The MHC-II^ΔEpi^ lungs also revealed reductions in Gata3^+^ T_RM_ cells (**Fig 5E**), suggesting dampened T_H_2 T_RM_ responses in LH lungs devoid of epithelial MHC-II. Consistent with T_H_2 cells being potent inducers of asthmatic airway remodeling ^1,3,19^, MHC-II^ΔEpi^ lungs exhibited milder airway hyperreactivity to methacholine compared to their MHC-II sufficient counterparts (**Fig 5G**). Notably, no genotype-dependent differences in cytokine secretion profiles of blood CD4^+^ T cells (**Fig 5F**), lung CD4^-^ T cells (**Fig S16D**) or lung non-T (CD3^-^) lymphocytes (**Fig S16E**) were observed. Thus, lung epithelial antigen presentation functions in a very tissue-specific (i.e., lung but not blood) and CD4^+^ T cell restricted-fashion. Taken together, epithelial cell antigen presentation skews CD4^+^ T_RM_ cells to T_H_1 and T_H_2 biology and regulates airway hyperreactivity during neutrophilic allergic airways disease.

**Figure 5:**
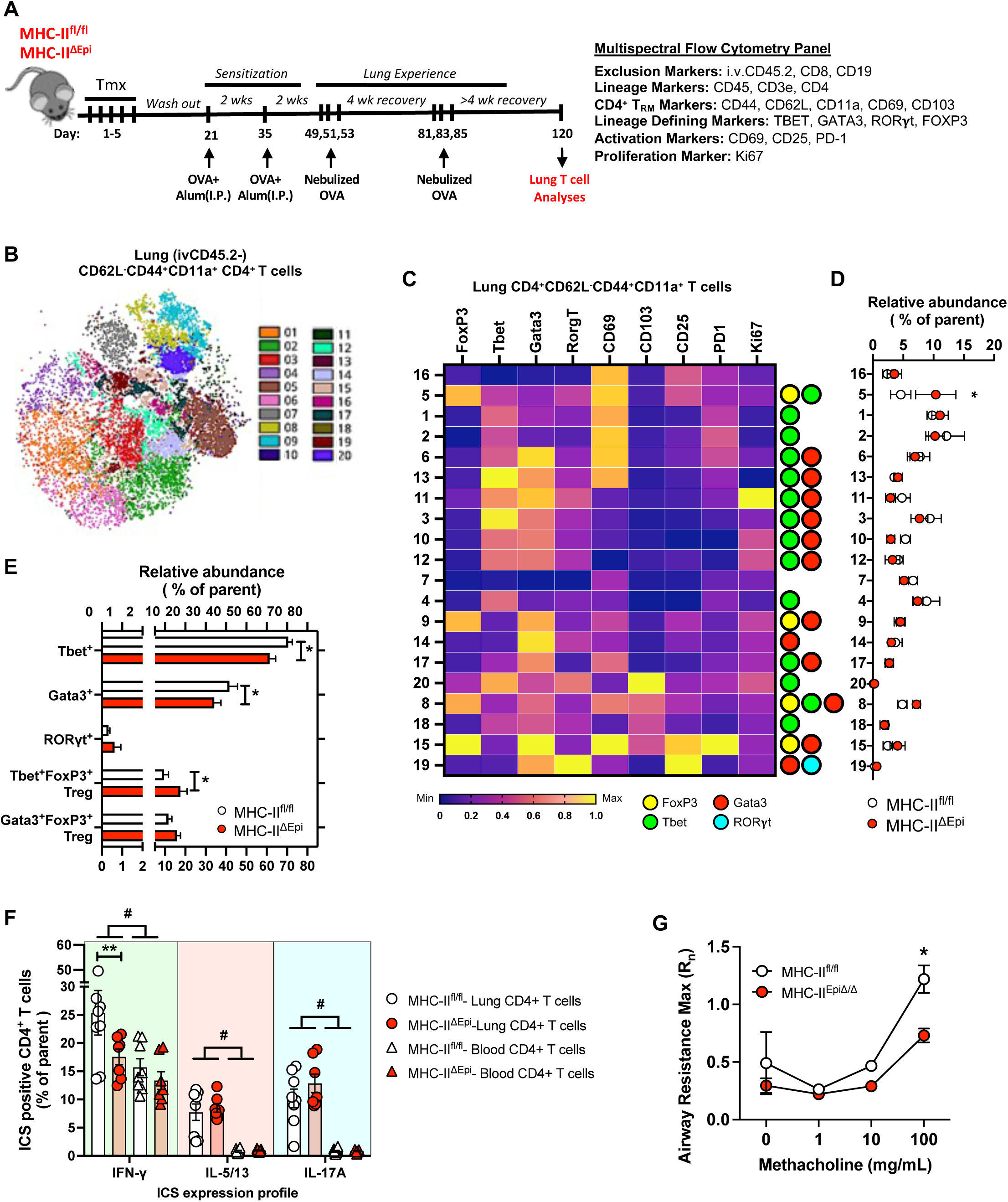
Antigen presentation by epithelial cells governs CD4^+^ T_RM_ cells in allergen-experienced lungs. **A)** Schematic of experimental timeline and antibody panel used. **B)** Phenograph clustering overlaid on opt-SNE projection depicting lung (ivCD45.2^-^) effector memory (CD62L^-^CD44^+^CD11a^+^) CD4^+^ T cells concatenated from MHC-II^fl/fl^ and MHC-II^ΔEpi^ LH lungs on day 120. **C)** Heat map depicting normalized expression levels of distinct molecules on lung effector memory CD4^+^ T cell clusters. Lineage determining transcription factor (LDTF) status of each cluster is depicted on the *right*. **D)** Relative abundance of each cluster of lung effector memory CD4^+^ T cells in MHC-II^fl/fl^ and MHC-II^ΔEpi^ LH mice at day 120. Two-way ANOVA with two-stage step-up method of Benjamini, Krieger, and Yekutieli to correct for multiple comparisons. FDR *q* value: *≤ 0.05. **E)** Cumulative frequencies for relative abundances of CD4^+^ T_RM_ cell positive for specified LDTF. Two-way ANOVA with Fisher’s LSD Test. **F)** Intracellular cytokine staining (ICS) profile of lung (i.v.CD45.2^−^) and blood (i.v.CD45.2^+^) CD4^+^ T cells isolated from MHC-II^fl/fl^ and MHC-II^ΔEpi^ LH mice on day 120 and stimulated with PMA/Ionomycin *ex vivo*. Two-way ANOVA with Fisher’s LSD Test. *p* value: ^#^≤ 0.05 comparison between lung and blood; *≤ 0.05 genotype-dependent comparison within lungs; ^Φ^ ≤ 0.05 genotype-dependent comparison within blood. **G)** Airway reactivity in response to increasing doses of methacholine in mice at baseline as measured by Flexivent. The measurements represent Maximum Rn values. Two-way ANOVA with Fisher’s LSD Test. *p*-value: *≤ 0.05, **≤ 0.01. All data have n≥5 mice, 2-3 experiments; mean ± SEM.

### Antigen presentation by epithelial cells regulates allergic airway neutrophilia

Altered phenotypes of lung T_RM_ cells due to epithelial MHC-II ablation led us to question how the absence of epithelial MHC-II affects the allergic neutrophilia response (**Fig 6A**). While mice of both genotypes possessed comparable BAL macrophage and lymphocyte numbers (**Fig S16F-H**), the inflamed MHC-II^ΔEpi^ mice showed greater neutrophil numbers within BAL **(Fig 6B)** and higher frequencies and numbers of extravascular (ivCD45.2^-^) neutrophils, but not eosinophils, in lungs after allergen challenge **(Fig 6C,D**). Furthermore, worsened neutrophilia was accompanied by elevated BAL CXCL5 levels (**Fig 6E**) and exacerbated airway edema **(Fig 6F),** despite comparable abundances of RORγt^negative/low^ T_H_17 T_RM_ cells (**Fig 6G)** and resulting IL-17A levels (**Fig S16I**) within MHC-II^ΔEpi^ lungs. Thus, lung epithelial MHC-II limits CXCL5 accumulation and neutrophilic inflammation in allergic airways by mechanisms distinct from, but seemingly downstream of T_H_17 T_RM_ cells.

**Figure 6:**
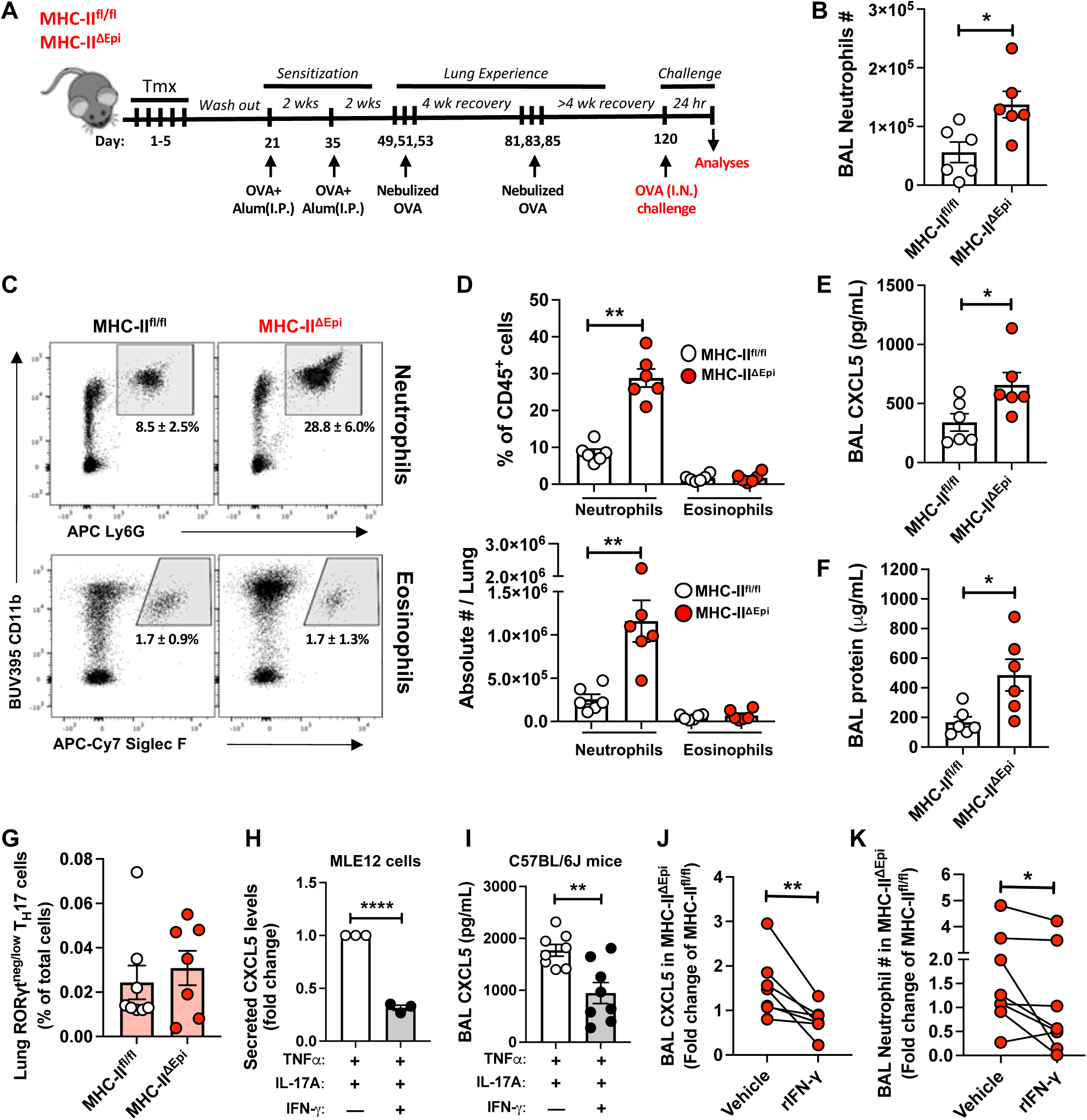
Lung epithelial MHC-II curtails severe allergic airways neutrophilia. **A)** Schematic of experimental timeline. **B)** Total neutrophil numbers in bronchoalveolar lavages (BAL) from MHC-II^fl/fl^ and MHC-II^ΔEpi^ LH mice 24 post OVA challenge. Mann-Whitney test. **C)** Representative dot plots depicting lung (ivCD45.2^-^) neutrophils and eosinophils as fraction (%) of CD45^+^ cells 24 hours post OVA challenge, mean ± SD. **D)** Numbers of lung (ivCD45.2^-^) neutrophils and eosinophils in MHC-II^fl/fl^ and MHC-II^ΔEpi^ LH mice 24 hours post OVA challenge. Mann-Whitney test. **E)** BAL CXCL5 in MHC-II^fl/fl^ and MHC-II^Δepi^ LH mice 24 hours post OVA challenge. Mann-Whitney test. **F)** Lung damage expressed as BAL protein content. Mann-Whitney test. **G)** Relative frequency of RORγt^negative/low^ IL-17A^+^ lung (ivCD45.2^-^) CD4^+^ T cells in MHC-II^fl/fl^ and MHC-II^ΔEpi^ LH mice. Mann-Whitney test. **H)** CXCL5 released by mouse lung epithelial (MLE12) cells treated with TNFα plus IL-17A with or without IFN-γ for 6 hours. Unpaired t test. **I)** BAL CXCL5 in C57BL/6J mice 7 hours post administration of specified cytokine cocktails. Mann-Whitney test. **J)** BAL CXCL5 and **K)** BAL neutrophil numbers in MHC-II^ΔEpi^ LH mice challenged with OVA plus vehicle or OVA plus IFN-γ, 8 hours post challenge. Data presented as fold change over cage-mate MHC-II^fl/fl^ LH mice. Mann-Whitney test. *p*-value: *≤ 0.05, **≤ 0.01. All data have n≥4 mice, 2 experiments; mean ± SEM.

### T_H_1 cytokine IFN-γ curbs CXCL5 and the excess airway neutrophilia of MHC-II^ΔEpi^ mice

Because T_H_1 biology and its effector cytokine IFN-γ can inhibit T_H_17 cell responses ^75^ and were curtailed by epithelial MHC-II deficiency (**Fig 5E,F**) alongside excess CXCL5 and neutrophils (**Fig 6C-E**), we asked whether IFN-γ might influence CXCL5 expression downstream of IL-17A signaling in the inflamed lung. Indeed, recombinant IFN-γ (rIFN-γ) blunted IL-17A-induced CXCL5 secretion by inflamed mouse lung epithelial cells *in vitro* (**Fig 6H**) and by mouse airways *in vivo* **(Fig 6I)**. Thus, IFN-γ from T_H_1 T_RM_ cells may prevent CXCL5 production and resultant neutrophilia driven by IL-17A in allergen experienced airways. Conversely, diminished T_H_1 numbers (and hence reduced IFN-γ) due to epithelial MHC-II deficiency may lead to uncontrolled CXCL5 production and exacerbate neutrophil recruitment. If so, the excessive neutrophilia phenotype due to epithelial MHC-II deletion should be corrected by supplementing IFN-γ. Consistent with this, exogenous rIFN-γ (concomitant with allergen challenge) reduced CXCL5 levels (**Fig 6J**) and neutrophil influx (**Fig 6K**) in airways of allergic MHC-II^ΔEpi^ mice. Of note, the Muc5ac^high^ airway secretory epithelial cells that are predominant producers of CXCL6 in adult human airways also express receptors for IFN-γ (**Fig S13G, S14H,** and **S15F**)^14^, suggesting responsiveness to this T_H_1 cytokine, and implicating this regulatory pathway in human lungs.

### IFN-γ inhibits allergic airway neutrophilia

The ability of recombinant IFN-γ to rescue excess neutrophilia in allergic MHC-II^ΔEpi^ mice suggested a finding of potential translational value, if such a strategy could mitigate allergic neutrophilia more broadly. Given the lack of effective therapies for patients with neutrophilic asthma, and the availability of rIFN-γ as an FDA-approved treatment for chronic granulomatous disease and severe malignant osteopetrosis ^76^, we explored IFN-γ as potential opportunity to mitigate neutrophilic allergic airways disease. **(i)** Consistent with the immunoregulatory role played by IFN-γ during allergic airway neutrophilia, IFN-γ knockout LH mice displayed unrestrained neutrophilic accumulation in the airways post OVA challenge when compared their No-LH counterparts (**Fig 7A-D**). **(ii)** We next asked if rIFN-γ delivered as a prophylactic (Px) prevented neutrophilic asthma exacerbations (**Fig 7E**, *blue track*). A single intranasal instillation of rIFN-γ delivered during the OVA challenge was sufficient to prevent allergic airway neutrophilia (**Fig 7F**) and resulting damage (**Fig 7G**) without affecting IL-17A levels in the LH lungs (**Fig S17A**). Although encouraging, asthma patients experience sudden and unpredictable onsets of allergic asthma exacerbation. (**iii**) Therefore, we also tested whether therapeutic (Tx) delivery of rIFN-γ (delivered 4 hours after inhaled OVA challenge, when sickness behavior due to acute asthma exacerbations are detectable in LH mice, **Fig S1E**) can limit severity of neutrophilic asthma (**Fig 7A**, *red track*). The systemic administration of rIFN-γ potently suppressed allergic airway neutrophilia (**Fig 7F**) and airway edema (**Fig 7G**) without affecting IL-17A in the LH lungs (**Fig S17A**) when delivered as a therapeutic regimen. These suppressive effect of rIFN-γ on airway neutrophilia can be retrospectively confirmed in humans, since nebulized or subcutaneously administered rIFN-γ robustly reduced BAL CXCL5 levels and neutrophils in airways of patients with idiopathic pulmonary fibrosis ^77^ or pulmonary tuberculosis^78^. Of note, despite favorable anti-neutrophilic outcomes, neither prophylactic nor therapeutic rIFN-γ delivery improved airway hyperreactivity in the acute setting of allergic asthma exacerbation tested in our studies (**Fig 7H**). However, since IFN-γ signaling blunts T_H_2 cell development and pathogenesis in asthma ^79–84^, it is plausible that repeated rIFN-γ therapy may gradually show positive outcomes with respect to airway physiology as well. Indeed, small independent clinical studies in patients with steroid-resistant asthma revealed that recurrent systemic administration of rIFN-γ improved lung function and reduced severity of asthma outcomes^85,86^. Thus, our findings suggest that IFN-γ (and the pathways it triggers in epithelial cells) could represent a rational promising approach to mitigating neutrophilic asthma.

**Figure 7:**
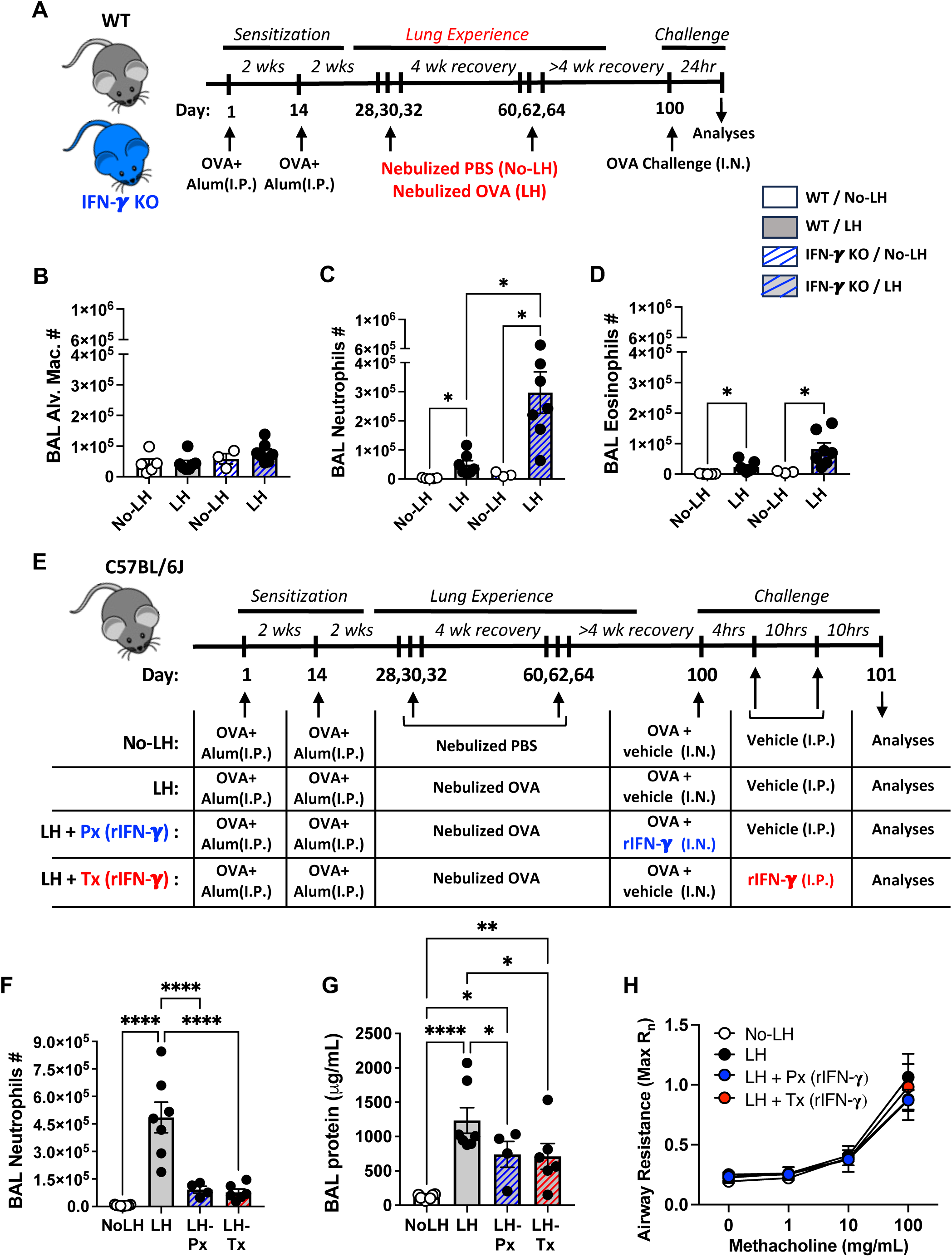
IFN-γ suppresses allergic airway neutrophilia. **A)** Schematic of experimental timeline. Total numbers of **B)** alveolar macrophages, **C)** neutrophils, and **D)** eosinophils in WT and IFN-γ knockout No-LH and LH mice 24 hours post OVA challenge. One-way ANOVA with Fisher’s LSD Test. **Note**: The Y axes in **Figs 7B-D** are adjusted to a similar scale to facilitate direct comparisons of airway inflammatory profiles. **E)** Schematic of experimental timeline. **F)** Total neutrophil numbers in BAL from mice 24 post OVA challenge with specified treatment modalities. One-way ANOVA with Fisher’s LSD Test. **G)** Lung damage expressed as BAL protein content. One-way ANOVA with Fisher’s LSD Test. **H)** Airway reactivity in response to increasing doses of methacholine in mice 24 hours post OVA challenge with specified treatment modalities as measured by Flexivent. The measurements represent Maximum Rn values. Two-way ANOVA with Fisher’s LSD Test. *p* value: *≤ 0.05, **≤ 0.01, ***≤ 0.001, ****≤ 0.0001. All data have n≥4 mice, 2-3 experiments, mean ± SEM.

## DISCUSSION

Recurrent but transient exposures of sensitized mice to inhaled ovalbumin seeds their lungs with a mature epithelial and immune ensemble that mirrors adult human lung biology, instills human predisposition to steroid-resistant and stable neutrophilic asthma, and thus provides a tractable animal model for this vexing disease. A notable feature of such experienced lungs includes establishment of diverse clusters of tissue-resident CD4^+^ T_RM_ cells comprised of unconventional RORγt^negative/low^ T_H_17 cells. Reactivation of these CD4^+^ T_RM_ cells leads to rapid secretion of IL-17A, inducing expression of CXCL5 by Muc5ac^high^ airway secretory epithelial cells which elicit peribronchial neutrophilic infiltration and severe disease. Antigen presentation by epithelial cells regulates disease severity by supporting protective T_H_1 T_RM_ cell activities, which curb production of CXCL5 and neutrophilia driven by IL-17A in allergen experienced airways. Thus, our studies identify key cellular and molecular constituents of biology relevant to human neutrophilic asthma, establishing a role for CD4^+^ T_RM_ cell-epithelial cell crosstalk in calibrating disease severity and identify IFN-γ (and immunoregulatory pathways it triggers) as a promising therapeutic avenue against the disease with direct translational implications.

Based on the age of diagnoses, asthma can be broadly classified into two broad phenotypes: childhood asthma which is dominated by T helper 2 (T_H_2) cell-driven steroid-sensitive eosinophilic inflammation and late-onset asthma associated with lung-damaging, steroid-refractory neutrophilic inflammation ^1,3,5,8^. A subset of patients transitions from treatment-responsive disease to become treatment-resistant over time ^9–12^. Pathways leading to these differential outcomes in an age-dependent manner need further understanding and study. Our work suggests that ‘T_H_2- high’ eosinophilic and ‘T_H_2- low’ neutrophilic endotypes of asthma may not be separate diseases but may represent extremities of a continual spectrum (**Fig S18A**). The progression in asthma endotypes may thus be linked with frequency and extent of aeroallergen exposure that occurs with time and advancing age. Specific elements of aeroallergenic experience may differentiate progression of eosinophilic asthma to neutrophilic disease from induction of antigenic tolerance, including the frequencies in durations of exposure ^49^.

Our studies identify a previously unrecognized diversity within the CD4^+^ T_RM_ cell pool residing in lungs prone to neutrophilic asthma. CD4^+^ T_RM_ cells within allergen experienced murine and human lungs have been reported to be of T_H_2- or Treg- T_RM_ cell phenotypes ^13,15–19,87^. The present studies extend roles for CD4^+^ T_RM_ cells to neutrophilic asthma, and show that allergic lungs can harbor diverse T_RM_ cell types which include Treg, T_H_1, T_H_2, and T_H_17 T_RM_ cells among a yet larger pool of T_RM_ lymphocytes that are negative for all the major LDTFs often tested. CD4^+^ T_RM_ cells in lungs with inhaled allergen history could produce IL-17A with minimal to no expression of the T_H_17-defining transcription factor RORγt. Although surprising, inspection of a scRNA-Seq studies profiling airway and peripheral blood CD4^+^ T cells of adult asthmatic and healthy humans confirmed the T cell heterogeneity observed in mice, including the unconventional RORγt^negative/low^ T_H_17 CD4^+^ T_RM_ cells ^14,70^, providing human disease correlates to our murine discoveries. Independent of the lungs or allergies, population of T_H_17 cells recently described as pathogenic in a mouse model of multiple sclerosis ^88^ also exhibited minimal RORγt expression compared to their homeostatic T_H_17 counterparts extending the significance of such unconventional T_H_17 cells to diverse allergic and autoimmune diseases of human health relevance. Further studies are warranted to determine whether these poorly defined pathogenic T_H_17 cells may be amenable to pharmacologic intervention.

Lung epithelial cells display anatomically-segregated abilities to communicate with CD4^+^ T cells and neutrophils during allergic airways neutrophilia (**Fig S18B**). Constitutive MHC-II expression on alveolar type 2 (AT2) epithelial cells has been reported before ^89,90^ and AT2 MHC-II was described to induce tolerance to inhaled allergens in a mouse model of eosinophilic asthma^71^. Whether epithelial antigen presentation may be involved in neutrophilic asthma was unclear. Here, we find Muc5ac^high^ secretory cells in the airways to be an especially immunorelevant cell type. This was reflected in their high expression of antigen-presentation molecules and simultaneous elaboration of CXCL5 (or CXCL6 in humans) within experienced airways. In the setting of pneumococcal pneumonia, T_H_17 T_RM_ cells, via secretion of IL-17A, instigate enhanced secretion of CXCL5 from lung epithelial cells to enforce rapid neutrophil recruitment and anti-bacterial immunity ^41,63^. Our current study extends this cell signaling axis to neutrophilic asthma and suggests that CXCL6 (the human ortholog of murine CXCL5) might be a good biomarker for neutrophilic asthma. CXCL6 correlates with disease severity in asthmatics ^91–93^. The observation that secretory cells are select sources of murine CXCL5 or human CXCL6 in neutrophilic allergic airways disease, in conjunction with a previous report demonstrating that depletion of airway secretory cells reduces T_H_2 cell responses, eotaxin production, and eosinophil recruitment during eosinophilic asthma ^94^, highlight these cells as consistently instrumental to both T cells and granulocytes in the allergic lung. CD45^-^EpCAM^-^ stromal cells were additional sources of CXCL5 in neutrophilic allergic airways, bolstering the expanding knowledge about immunological functions for stromal cells during lung diseases ^95–99^. Which cell types within the structural cell fraction are CXCL5 producers is unclear but may include peribronchial fibroblasts, airway smooth muscle cells, pericytes, and/or endothelial cells ^73,100–104^.

Epithelial antigen presentation regulates severity of allergic airway neutrophilia by instructing CD4^+^ T_RM_ cell activities. While epithelial MHC-II supported formation of T_H_2 T_RM_ cells and bolstered airway hyper-reactivity, it also enriched the lung T_RM_ pool for T_H_1 T_RM_ cells that appeared to curb allergic airway neutrophilia. Thus, lung epithelial antigen presentation functions as a rheostat that sets the fine balance between two clinical symptoms of neutrophilic asthma, airway hyper-reactivity and allergic airway neutrophilia. It does so by altering the balance of T_RM_ cells and reinforcing protective T_H_1 cell activities in the airways (**Fig S18B**). Elevated T_H_1 cell numbers are observed in asthmatic lungs ^105,106^ and may reflect an attempt of the inflamed airways to limit allergic inflammation, implicating T_H_1 cells as mitigating agents rather than pathogenic drivers in this allergic disease. Given that MHC-II on epithelial cells is key to airway T_RM_ cell biology, the polymorphisms in MHC-II related genes that have been linked with adult-onset asthma ^107,108^ may be significant for influencing the relative frequencies and activities of CD4^+^ T_RM_ cell lineages in the airways.

Expanding beyond its established roles in stifling T_H_2-high eosinophilic asthma ^79–84^, we now surmise that T_H_1 cytokine IFN-γ can regulate pathogenic T_H_17-driven neutrophilic asthma. However, rather than via the previously reported direct inhibition of T_H_17 cell differentiation and activities by IFN-γ ^75^, the protective function of T_H_1 activities during neutrophilic asthma are exerted downstream of the T_H_17 cells, by IFN-γ blunting IL-17A-induced CXCL5 production in epithelial cells of the allergic airways. This defines a novel pathway for regulation of type 17 inflammation by IFN-γ without direct inhibition of T_H_17 cells. Whether IFN-γ suppresses *de novo* CXCL5 transcription or uncouples the IL-17A mediated stabilization of CXCL5 mRNA is under active investigation ^41,109^. Nevertheless, our discovery of the suppressive effects of IFN-γ on airway neutrophilia and lung damage suggest this T_H_1 effector cytokine and the pathways its triggers as promising avenues for future research with potential translational and clinical utility for neutrophilic asthma. This is also supported by an empirical clinical study ^85^.

Taken together, our studies using an ovalbumin induced mouse model of allergic airways disease suggest that asthma endotypes may represent stages in a continuum of disease progression and offer insights into mechanisms underlying pathophysiology of neutrophilic asthma. Our results implicate a T_H_17-CXCL5 axis in allergic airway neutrophilia, with IL-17A in such lungs coming from unusual RORγt^negative/low^ CD4^+^ T cells. We define lung epithelial cells as key regulators and pathogenic effectors in the immune circuitry programmed within allergen-experienced airways of mice and humans, owing to their ability to instruct CD4^+^ T_RM_ cell activities (via antigen presentation) and to direct airway neutrophilia (by acting as signaling nodes that integrate cues from pathogenic T_H_17 T_RM_ cells and T_H_1 T_RM_ cells to effectively fine-tune CXCL5 release). These studies advance the concept that IFN-γ can curb effects of IL-17A on epithelial cells (including specifically their CXCL5 expression), leading to studies which now suggest that IFN-γ and/or the pathways this cytokine triggers deserve further investigation as promising therapeutic avenues for mitigating neutrophilic asthma.

## LIMITATIONS OF THE STUDY

To focus selectively on how inhaled allergen experience may remodel the lung, we chose OVA instead of a clinically relevant but complex and less-defined allergen preparations of house-dust mites (HDM), cockroaches, or fungi. This was in order to minimize contributions of confounding factors like trained immunity (or other off-target immune conditioning events) and to allow precise and effective determination of the effects of inhaled allergen experiences on disease outcomes in sensitized hosts. This however limits translation because OVA is not an asthma-relevant allergen. Also, the mice in our study were initially sensitized to OVA with alum as an adjuvant via the peritoneum, which is not akin to the human experience where allergen exposure mostly occurs at the mucosal site. Thus, the first exposure of T cell-instructive dendritic cells to allergen occured via peritoneal DCs rather than those from the airways, which may have different phenotypes. Finally, while rIFN-γ treatment was sufficient to reduce severity of allergic airways disease in our reductionist mouse model as well as some steroid-resistant human asthmatics^85,86^, T_H_1 cells and IFN-γ can drive neutrophilic airway inflammation in other conditions^110–112^. Thus, we emphasize caution and the need for further preclinical studies before considering IFN-γ as a potential means of limiting neutrophilic asthma. Instead, our findings direct us towards exploring the signaling pathways triggered by IFN-γ in this and other preclinical models of asthma to see if translational and clinical opportunities against neutrophilic asthma may arise by studying this immunoregulatory biology more carefully.

## MATERIALS AND METHODS

### LEAD CONTACT AND MATERIALS AVAILABILITY

Correspondence and requests for information, resources and reagents should be directed to Anukul T. Shenoy. All data will be made available by the corresponding authors upon reasonable request.

#### Mice

6-week-old C57BL/6J (Stock# 000664), IL-17A/F knockout (B6.Cg-Il17a/Il17f^tm1.1Impr^ Thy1^a^/J, Stock # 034140)^113^, and IFN-γ knockout (B6.129S7-Ifng^tm1Ts^/J, Stock# 002287)^114^ mice were obtained from Jackson labs (USA). SPC-GFP mice and Nkx2.1^cre/ERT2^H2-Ab1^fl/fl^ mice (henceforth called MHCII^ΔEpi^ mice) are described elsewhere ^69,71^ and were bred in-house. All breeders were maintained as homozygous floxed for H2-Ab1. For experiments all mice were homozygous for loxP sites flanking H2-Ab1 exon1 and were identified as Cre-positive or negative based on presence or absence of Nkx2.1^cre/ERT2^. Cage and littermate controls of both sexes were used for studies. 7-14-week-old mice were used for experiments and animals were housed in specific pathogen free environment on a 12-hour light-dark cycle, with ad libitum access to standard chow and water. Mice were euthanized using isoflurane overdose and death confirmed using pneumothorax before organ collections. All animal procedures were approved by the Institutional Animal Care and Use Committee at the University of Michigan at Ann Arbor and Boston University.

#### Experimental allergic airways eosinophilic disease

OVA (Sigma Aldrich) or sterile PBS was adsorbed on Alum (Sigma Aldrich) for 30 mins on shaker before sensitization of mice to OVA at dose of 25μg/100μL via intraperitoneal (i.p.) injection of 100μL on days 1 and 14. OVA or PBS sensitized mice were then exposed to nebulized solution of 2% OVA in sterile PBS for 30 mins in a nebulization chamber on days 28, 29 and 30. On day 31, mice were anaesthetized with ketamine and xylazine before receiving intranasal (I.N.) challenge with 100μL of 0.2% OVA in sterile PBS.

#### Experimental allergic airways neutrophilic disease

OVA (Sigma Aldrich) or sterile PBS was adsorbed on Alum (Sigma Aldrich) for 30 mins on shaker before sensitization of mice to OVA at dose of 25μg/100μL via intraperitoneal (i.p.) injection of 100μL on days 1 and 14. OVA sensitized mice were then split into two groups andexposed to nebulized solution of 2% OVA in sterile PBS or plain sterile PBS (for control mice) for 30 mins in a nebulization chamber on days 28, 30, 32 and days 60, 62, 64 with a 4 week recovery period in between. After at least 4 weeks or more recovery period, mice were then anaesthesized with ketamine and xylazine before receiving intranasal (I.N.) challenge with 100μL of 0.2% OVA in sterile PBS.

#### Pneumococcus infections

*Streptococcus pneumoniae* (*Spn*)-specific lung-resident CD4^+^ T_RM_ cells were generated as previously described ^41,63^. Briefly, mice were intratracheally infected with ∼10^6^ CFU of serotype 19F *Spn* (Strain EF3030) suspended in sterile saline on days 0 and 7 followed by recovery for 28–35 days. For pneumococcal infections to test antigen specificity of airway neutrophilia, mice were intratracheally infected with ∼10^6^ CFU of serotype 3 *Spn* (Sp3, ATCC 6303) suspended in sterile saline as previously described ^41^.

#### Physiological measurements of airway hyper-reactivity

For airway resistance assay, mice were injected with xylazine (10 mg/kg body weight), pentobarbital (100 mg/kg body weight) and pancuronium (0.5 mg/kg body weight), intubated and placed on a mechanical ventilator (Legacy flexiVent, SCIREQ). Ventilation was at 300 breaths/min (tidal volume 6–7 mL/kg body weight). Airway resistance was measured after airway delivery of nebulized methacholine in PBS (0, 1, 10, and 100 mg/mL). The plotted measurements represent Maximum Newtonian resistance (Rn) values.

#### Tamoxifen, Dexamethasone, antibody administration

Tamoxifen (Sigma) was dissolved in corn oil (Sigma) to 20mg/mL stock concentration and stored at 4°C. Mice were i.p. injected at 100mg/kg of body weight for 5 consecutive days followed by a washout of at least 2 weeks before experimentation. Dexamethasone (1mg/kg body weight) was delivered intraperitoneally 72 and 24 hours prior to the OVA challenge to assess steroid responsiveness of the mice. CD4^+^ T_RM_ cells were depleted as described previously^41^. Briefly, mice were administered 500μg and 100μg of GK1.5 (BioXcell, West Lebanon, NH) i.p. and i.n. respectively, both 72 and 24 hours prior to the OVA challenge.

#### Lung Histology

Euthanized mice were exsanguinated, and their tracheas cannulated with 25 gauge butterfly needle before inflation of lungs with 4% paraformaldehyde at 23 cm H_2_O pressure. The left lobes were paraffin embedded after overnight fixation in 4% paraformaldehyde, cut into 5μm thin sections and stained using hematoxylin and eosin.

#### Bronchoalveolar lavage (BAL) collection and analyses

Euthanized mice were exsanguinated, and their tracheas cannulated with a 18 gauge cannula followed by 6 rounds of lavage with sterile PBS. The supernatant from first lavage was stored at -80°C for BAL chemokine and protein measurements while the cell pellets from all lavages per mouse were compiled to perform flowcytometry or cytospins, differential staining and airway cellularity enumeration. BAL proteins levels were measured using standard BCA assay. BAL CXCL1, CXCL2, CXCL5 and CXCL10 levels were measured using ELISA kits from R&D Systems using the manufacturer’s protocols.

#### Lung digestion for flow cytometry

To enumerate of extravascular versus intravascular fraction of lung CD4^+^ T cells and myeloid cells, anesthetized mice were retro-orbitally administered 2μg anti-CD45.2 antibody 3-5 minutes prior to euthanasia ^115^. Lungs were collected in RPMI 1640 with 10% FBS for flow cytometry. Single-cell suspensions were prepared by digestion of lungs as previously described by *Smith et al.* ^63^. For high throughput flow cytometry and FACS sorting of lung epithelial cells, single cell suspensions of lungs were generated as previously described by *Shenoy et al.* ^69^. Cells in single cell suspensions were blocked with TruStain αCD16/CD32 Fc-Block (BioLegend). Flow cytometry was performed on LSR Fortessa or LSR II Flow Cytometer (both BD Biosciences) and data was analyzed with FlowJo software (BD Biosciences). High-dimensional multi-parameter spectral flow cytometry was performed on Aurora (Cytek). SpectraFlo (Cytek) software was utilized for spectral unmixing of the data using ordinary least square algorithm and and data were analyzed using FlowJo (BD Biosciences). Gating strategies are provided in the Supplemental figures and were based on use of Fluorescence minus one (FMO) controls.

For RNA-profiling, leukocytic (CD45^+^), epithelial (CD45^-^EpCAM^+^) and stromal (CD45^-^ EpCAM^-^) cell fractions from stained single cell suspensions were sorted into RLT buffer with beta-mercaptoethanol on ice using FACS-Aria II SORP (BD Biosciences) before proceeding to RNA extraction.

#### *Ex vivo* stimulation of CD4^+^ T cells

To allow identification of extravascular versus intravascular fraction of lung CD4^+^ T cells, anesthetized mice were retro-orbitally administered 2μg anti-CD45 antibody 3-5 min prior to euthanasia. Single cell suspension of lung leukocytes was prepared as described before and 2 × 10^6^ cells were stimulated *ex vivo* in 12 well plate with 250ng/mL Phorbol Myristate Acetate (PMA)(LC Laboratories, Woburn, MA) and 1.5μg/mL Ionomycin (Sigma, St. Louis, MO) in T cell stimulation media for 1hour at 37°C and 5% CO_2_. Monensin (Biolegend, Cat# 420701) and Brefeldin A (Biolegend, Cat# 420601) both at 1X final concentration were added to the cell suspension for last 5 hours at 37°C and 5% CO_2_. Cells were then processed for intracellular cytokine and transcription factor staining, as per manufacturer’s protocols.

#### Intracellular cytokine, protein and transcription factor staining for flow cytometry

For intracellular staining of transcription factors and Ki67, eBioscience Foxp3/ Transcription Factor Staining Buffer Set (Cat# 00-5523-00) was used as per manufacturer’s protocols. For intracellular cytokine staining, eBioscience Intracellular Fixation & Permeabilization Buffer Set (Cat# 88-8824-00) was used as per manufacturer’s protocols. For intracellular costaining of cytokines and transcription factors, eBioscience^TM^ Foxp3/ Transcription Factor Staining Buffer Set (Cat# 00-5523-00) was used.

#### Algorithmic analysis of single cell fluorescence datasets

Data processing pipeline was established using the Omiq.ai cloud computation platform (Omiq) as previously described ^69^. Briefly, live CD45^+^CD4^+^CD19^-^CD45.2^+^ for lung intravascular (‘blood’) cells or live CD45^+^CD4^+^CD19^-^CD45.2^-^ for lung extravascular (‘lung’) T cells were concatenated, asinh transformed (cofactor = 6000) and clustered with Phenograph algorithm (k=20, distance metric = euclidean) ^67,68^ followed by prejection in opt-SNE space (perplexity=30, theta=0.5, opt-SNE endpoint = 5000; PCA pre-initialization embedding). For data in Figure 5, CD62L^-^ CD44^+^CD11a^+^ clusters (as determined based on the MFI cutoff) were subsampled as memory T cell data. These data were then re-clustered with Phenograph (k=20, distance metric = euclidean) and projected into opt-SNE space (perplexity=30, theta=0.5, opt-SNE endpoint = 5000; PCA pre-initialization embedding). Clusters were color-coded and overlaid on the opt-SNE projections. Each marker MFIs of Phenograph clustered datasets were organized into hierarchically clustered heatmaps. Clusters were classified as positive for specific lineage-defining transcription factors (LDTF) based on the MFI value cut offs set by measuring the MFIs of LDTF-negative non-T cell populations sampled from the same dataset. This approach was preferred over the FMO-based cutoff calculation to alleviate the MFI difference caused by non-specific binding of anti-LDTF antibodies to cells that do not express them. Frequencies of each cell type and MFI values for individual markers were calculated from corresponding clusters per each animal and data were plotted and compared using Prism 8.0 (Graphpad).

#### Analyses of independently generated adult human lung single cell RNA-Seq datasets

Previously published single cell RNA-Seq datasets profiling epithelial cells and CD4^+^ T cells from blood and lungs of adult humans and asthmatics were interrogated for specified target transcripts using their respective interactive webtools ^14,73^. For reanalyses of human airways dataset of adult asthmatic and healthy controls from Alladina Sci. Immunol. 2023^70^, processed data was obtained from the GEO database (under accession number GSE193816). UMAP plots were generated using the *pl.umap* function in Scanpy^116^(version 1.9.4). Dot plot was created using *pl.scatter*.

Threshold lines were defined based on the mean gene expression values. For reanalyses of murine allergic airway eosinophilia dataset from Tibbitt et al. Immunity 2019^17^, raw data was downloaded from the GEO database (under accession number GSE131935). The raw count matrices were processed using Scanpy^116^ (version 1.9.4), following a standard pipeline: *pp.normalize_total, pp.log1p, pp.highly_variable_genes, tl.pca, pp.neighbors, tl.leiden,* and *tl.umap*, all with default parameters. UMAP plots were generated using the *pl.umap* function, and dot plot was created with *pl.scatter*. As with the human dataset, threshold lines were set based on the mean gene expression values.

#### Immunofluorescent staining

The trachea of euthanized mice were cannulated with an 18-gauge angiocath and 1.4 mL of Tissue-Tek Optimal Cutting Temperature (O.C.T.) compound (Sakura Finetek) was slowly instilled in the lung. Once the lungs were inflated, the left bronchus was tied using a suture, the lungs washed in sterile HBSS before embedding in cryomolds with O.C.T. and flash freezing at −80 °C until sectioning. Frozen 8 μm thin coronal sections of the left lungs that contained the whole face of the lungs including the entire airway tree structure were collected for further analyses. Sections with folds and/or tears were rejected. The sections that met our inclusion criteria were first fixed, washed, and permeabilized (with 0.2% Triton X) before blocking with Blocking buffer (PBS with 10% normal donkey serum and 3% BSA) followed by overnight incubation with rabbit antimouse CD4 (abcam, Cat# ab183685) and rat antimouse Ly-6G (clone 1A8; BD Biosciences) at 4 °C in a humidified chamber. Next day, sections were vigorously washed before incubation with Alexa 594 conjugated Affinipure donkey antirabbit IgG and Alexa 488 conjugated Affinipure donkey antirat IgG (Jackson Immunoresearch) at room temperature for 1 hr in a dark humidified chamber. All slides were counterstained with DAPI (Molecular Probes by Life Technologies, R37606) before mounting the sections with FluorSave (Millipore Calbiochem: 345789) and covering with coverslip for visualization. Of note, “no primary antibody control” were used to identify true events. Images were captured using a Leica DM4 microscope equipped with Leica DFC 7000T camera and processed using ImageJ 2.0.0-rc-69.

#### In vitro studies on lung epithelial cell line

Mouse lung epithelial (MLE12) cells were treated with cell culture media containing TNF-α (25ng/mL) + IL-17A (50ng/mL) with or without IFN-γ (50ng/mL) for 6 hours before collection of supernatant for CXCL5 ELISA. All mouse cytokines were purchased from R&D Systems.

#### RNA extraction and Real-time PCR

RNA was extracted from sorted cells using RNAeasy Micro Kit (Cat# 74004) as per manufacturer’s protocols and stored at -80°C. qRT-PCR was performed using the RNA-to-Ct kit (Life Technologies). Commercially available TaqMan gene expression assay primers and probes for cxcl1, cxcl2, cxcl5, cxcl10, muc5ac and 18S rRNA (Cat# 4319413E) from Applied Biosystems were used. The quantity of the detectable mRNA was calculated by normalizing to 18S rRNA from the respective sample and then expressed as fold change over mRNA levels of whole lung of naive mice.

#### *In vivo* recombinant IFN-γ administration

Mice were administered 100ng of recombinant mouse IFN-γ (or vehicle) in 100μL volume for intranasal Px-rIFN-γ administrations and 200ng of recombinant mouse IFN-γ (or vehicle) in 200μL volume for intraperitoneal Tx-rIFN-γ administrations at designated timepoints. Recombinant mouse IFN-γ was purchased from R&D Systems.

#### Statistical Analyses

Statistical analyses were performed using Prism 10 (Version 10.3.0, GraphPad). Differences were deemed statistically significant if the *p* value or FDR *q* value was ≤0.05. Each figure legend communicates the number of mice used, the experiment replicates performed and the statistical tests used to make comparisons. For all figures, data are represented as mean ± SEM, except for **Fig 1G and 6C** which depicts flowcytometry plots with data represented as mean ± SD).

## LIST OF SUPPLEMENTARY MATERIALS

Supplementary Figures **S1-S18** and Supplementary Figure legends.

## ACKNOWLEDGEMENTS

All graphics were created with BioRender.com. We thank Drs. George O’Connor, Frederic Little and John Bernardo for helpful discussions. We thank Olivia Harlow and Fang Ke for technical assistance, and the Dept. of Microbiology and Immunology at the University of Michigan for support with flowcytometry. We thank Brian Tilton and the Boston University Chobanian & Avedisian School of Medicine Flow Cytometry Core Facility (BU-FCCF) for assistance with FACS sorting. This work was supported by NIH grants including F32 HL147461 and K99 HL159258 to F.T.K., F31 HL142199 to K.A.B., R01 HL136725 to M.R.J. and A.F., R01 HG010883 to J.D.W., R01 GM120060 and R01 HL111449 to L.J.Q., R01 AI115053, R35 HL135756, and R33 HL137081 to J.P.M., and K99 HL157555, R00 HL157555 to A.T.S., plus T32 HL007035 for support of trainees. This work was also supported by Institutional funding and Endowment for Basic Science from University of Michigan to A.T.S.

## AUTHOR CONTRIBUTIONS

Conceptualization, A.T.S.; Experimental Design and Interpretation, V.R.R., F.C., A.C.B. J.P.M., and A.T.S.; Investigation, V.R.R., F.T.K., C.L.D.A., L.L., F-Z.S., E.N.N., C.V.O., K.A.B., A.R., W.N.G., I.M.C.M., C.H., and A.T.S.; Validation, A.C.B. and A.T.S.; Supervision, A.T.S.; Project Management, A.T.S.; Software, V.R.R., L.L., F-Z.S., A.C.B., and A.T.S.; Formal Analysis, V.R.R., L.L., A.C.B., and A.T.S.; Data Curation, V.R.R., F.C., A.C.B., and A.T.S.; Visualization, V.R.R., L.L., A.C.B., J.P.M., and A.T.S.; Resources, L.J.Q., M.R.J., A.F., J.D.W., F.C., A.C.B., J.P.M., A.T.S.; Writing-Original Draft, A.T.S.; Writing-Review & Editing, V.R.R., F.T.K., C.L.D.A., L.L. F-Z.S., E.N.N., C.V.O., K.A.B., A.R., W.N.G., I.M.C.M., C.T.H., L.J.Q., M.R.J., A.F., J.D.W., F.C, A.C.B., J.P.M., and A.T.S.

## DECLARATION OF INTERESTS

The authors declare no competing interests.

## SUPPLEMENTAL FIGURES AND FIGURE LEGENDS

**Figure S1:**
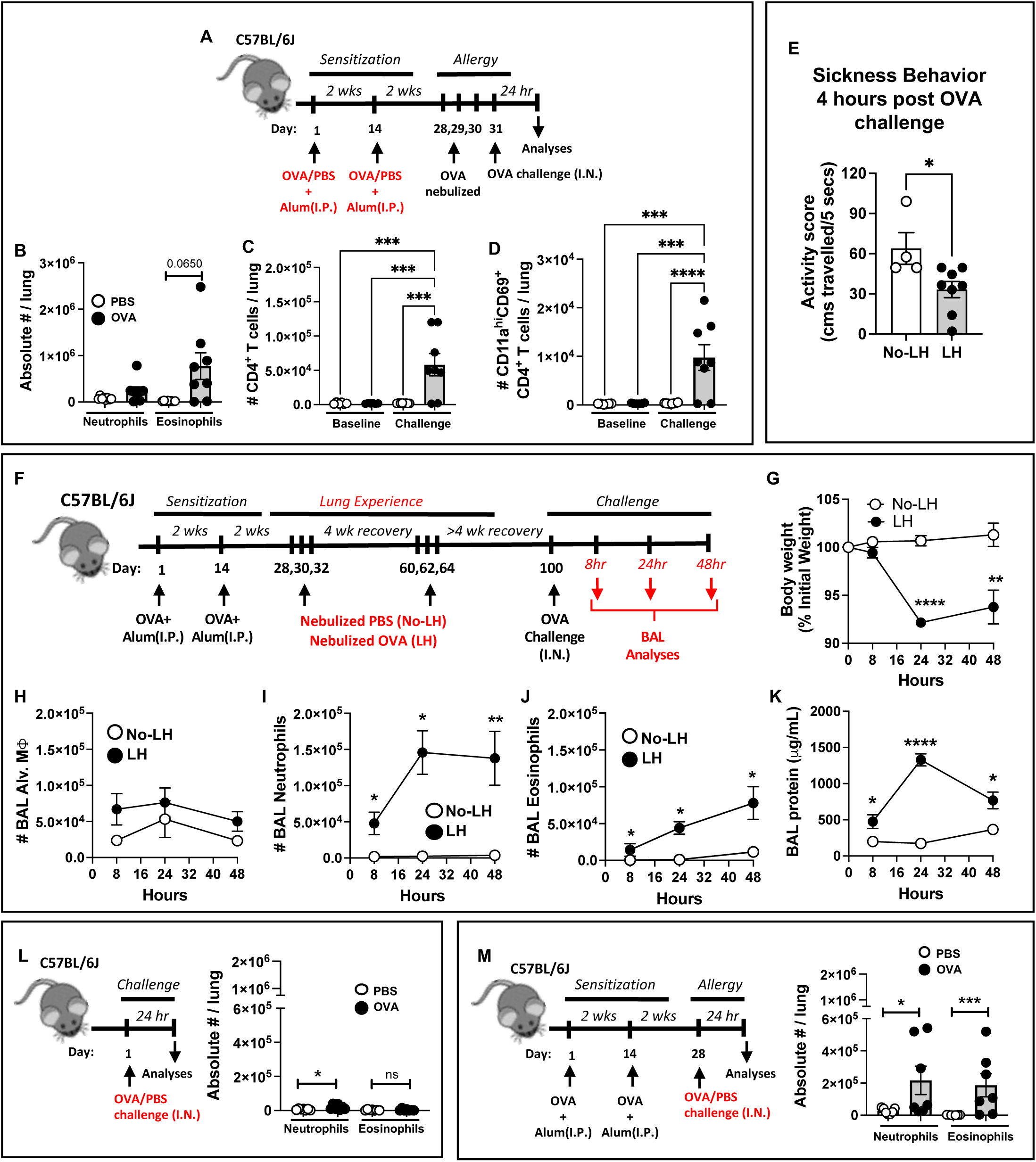
Comparisons of mouse models of OVA-induced eosinophilic and neutrophilic asthma. **A)** Schematic of experimental timeline used. **B)** Numbers of lung (ivCD45.2^-^) neutrophils and eosinophils 24 hours post final OVA challenge in OVA or PBS sensitized C57BL6/J mice. Mann-Whitney test. **C-D)** Numbers of **C)** lung (ivCD45.2^-^) CD4^+^ T cells and **D)** lung (ivCD45.2^-^) CD11a^hi^CD69^+^ CD4^+^ T cells at baseline and 24 hours post OVA challenge. One-way ANOVA with Fisher’s LSD Test. **E)** Activity score of LH and No-LH mice measured as distance travelled in cms within 5 seconds when assessed 4 hours post OVA challenge. Mann-Whitney test. **F)** Schematic of experimental timeline used. **G)** Body weight of of LH and No-LH mice at designated timepoint. Mann-Whitney test comparing LH to No-LH at each timepoint. Total number of **H)** alveolar macrophages, **I)** neutrophils, and **J)** eosinophils in bronchoalveolar lavages (BAL) from LH and NoLH mice at designated timepoints after OVA challenge. Mann-Whitney test comparing LH and No-LH mice at each timepoint. **Note**: The Y axes in **Figs S1H-J** are adjusted to a similar scale to facilitate direct comparisons of airway inflammatory profiles at each timepoints. **K)** Lung damage expressed as BAL protein content. Unpaired t test. **L)** Schematic of experimental timeline used and numbers of lung (ivCD45.2^-^) neutrophils and eosinophils 24 hours post PBS or OVA challenge in naïve C57BL6/J mice. Mann-Whitney test. **M)** Schematic of experimental timeline used and numbers of lung (ivCD45.2^-^) neutrophils and eosinophils 24 hours post PBS or OVA challenge in OVA sensitized C57BL6/J mice. Mann-Whitney test. *p* value: *≤ 0.05, **≤ 0.01. All data have n≥3 mice, 2 experiments, mean ± SEM. Note: The Y axes in **Fig S1L** and **Fig S1M** are adjusted to be similar with **Fig 1H** to facilitate direct comparisons.

**Figure S2:**
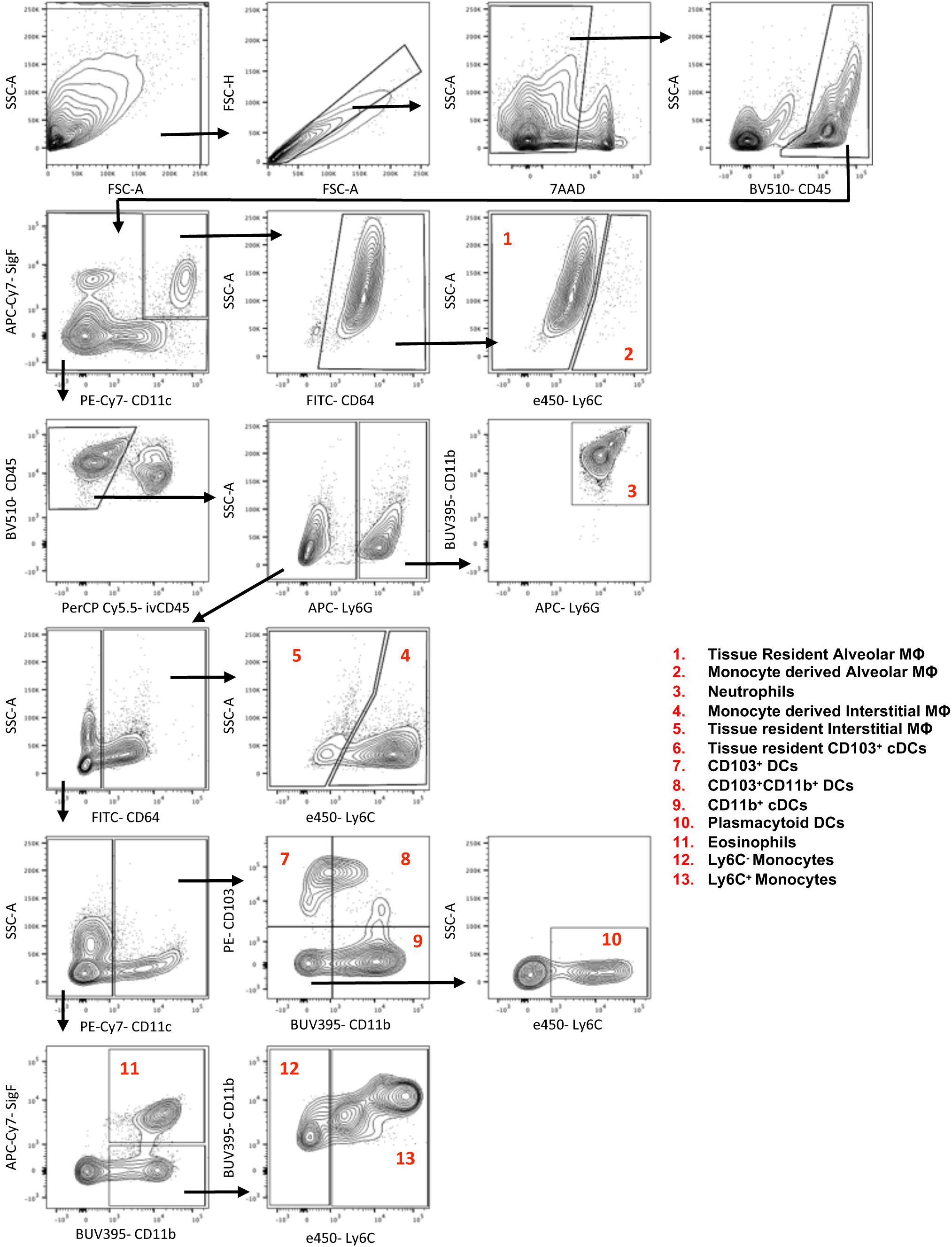
Gating strategy for comprehensive profiling of myeloid cells.

**Figure S3:**
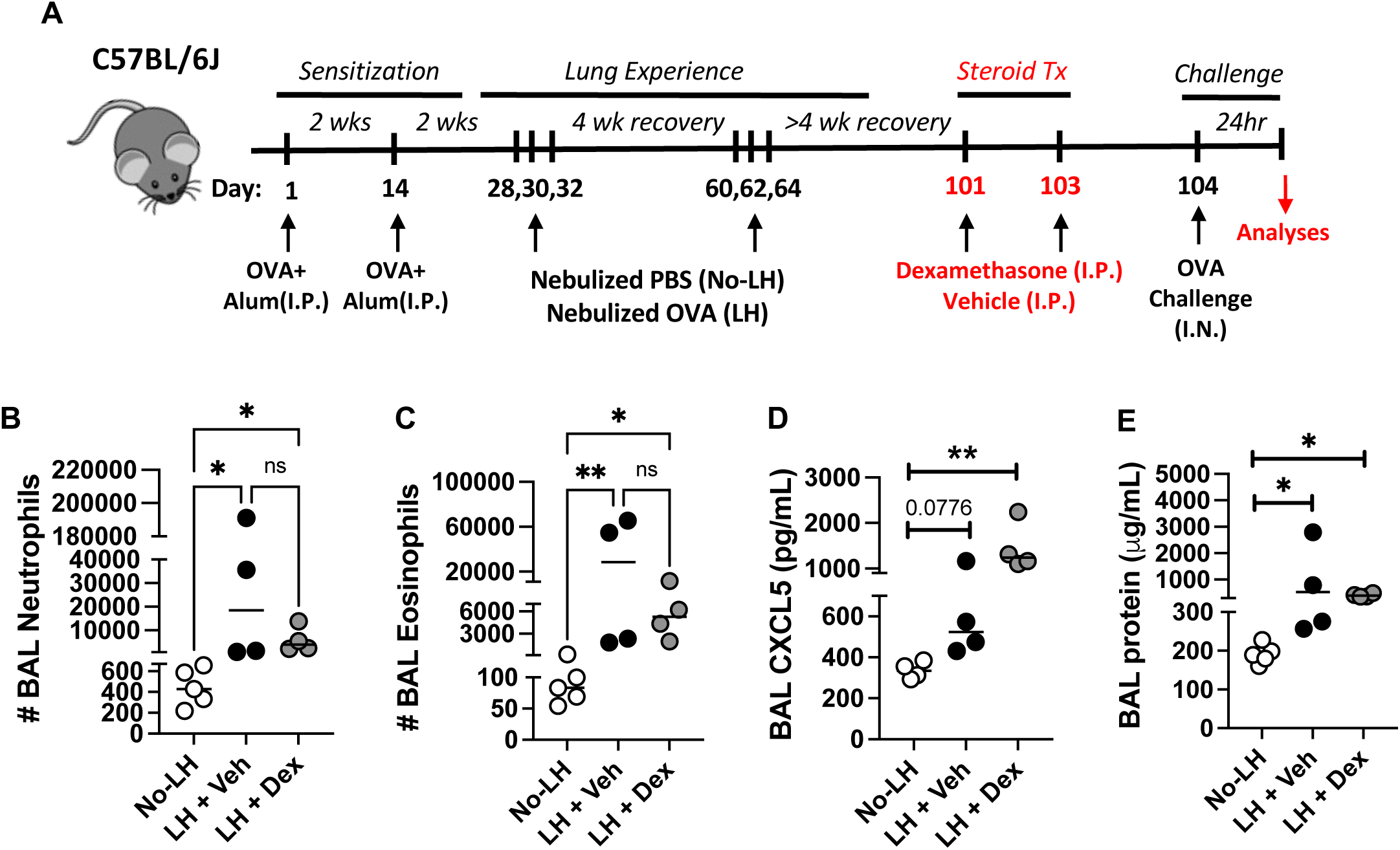
LH mice phenocopy steroid-resistant allergic airway neutrophilia. **A)** Schematic of experimental timeline used. LH mice were prophylactically administered high doses of dexamethasone (1mg/kg) or vehicle intraperitoneally 72- AND 24- hours- before OVA challenge for assessment of steroid resistance for allergic airway neutrophilia. Total numbers of **B)** neutrophils and **C)** eosinophils in bronchoalveolar lavages (BAL) from mice 24 hours post OVA challenge. Levels of **D)** BAL CXCL5 and **C)** lung damage in mice 24 hours post OVA challenge. All data have n≥4 mice, 2 experiments, mean ± SEM, Kruskal-Wallis test, *p* value: *≤ 0.05, **≤ 0.01.

**Figure S4:**
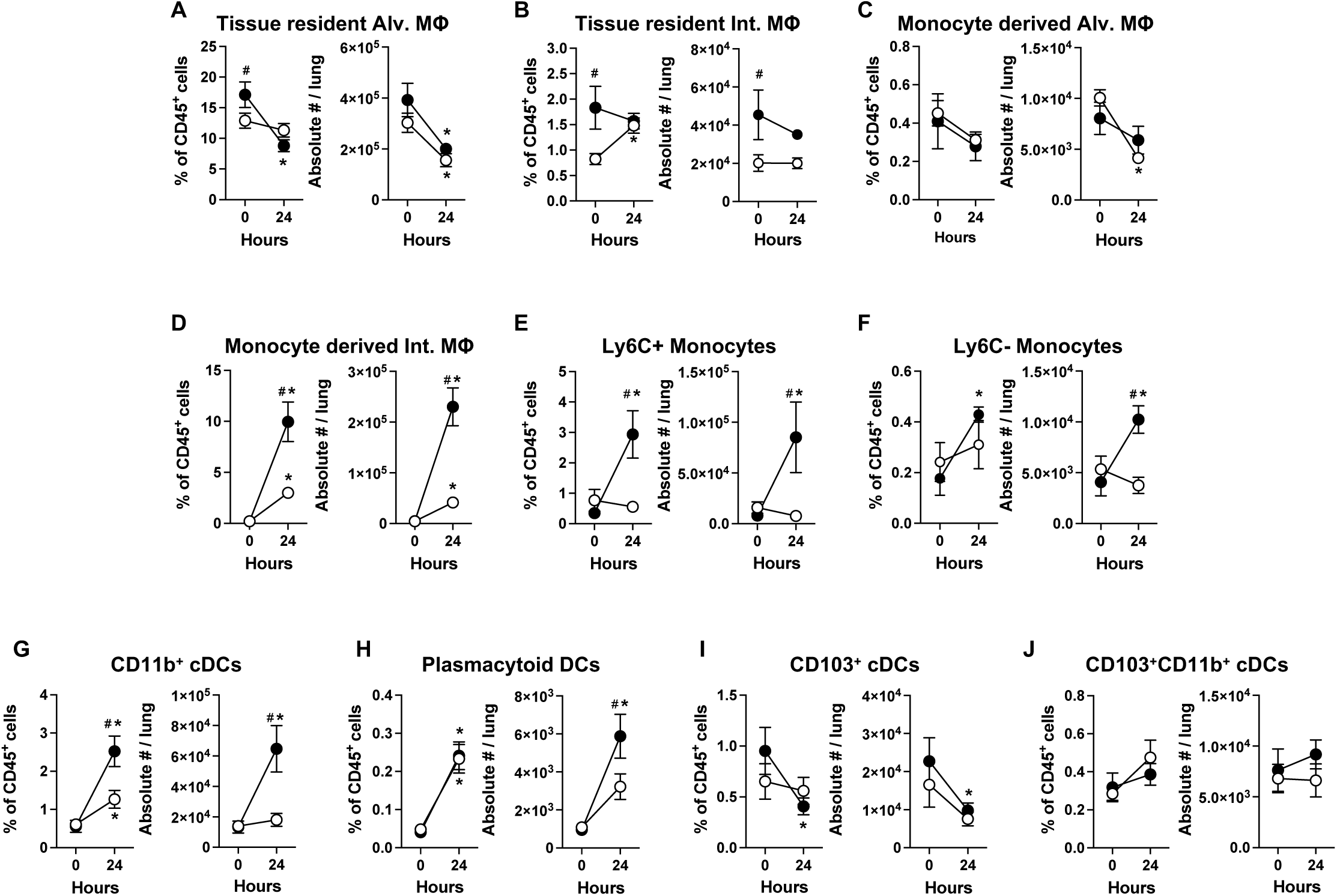
LH mice display extensive remodeling of lung myeloid landscape. Numbers of distinct lung (ivCD45.2^-^) myeloid cells at baseline and 24 hours post OVA challenge in no-LH (white) and LH mice (black), Two-way ANOVA with Fisher’s LSD Test. *p* value: *≤ 0.05 for comparisons within same group across time-points and ^#^≤ 0.05 for comparisons between groups at same time-point. All data have n≥5 mice, 2 experiments, mean ± SEM

**Figure S5:**
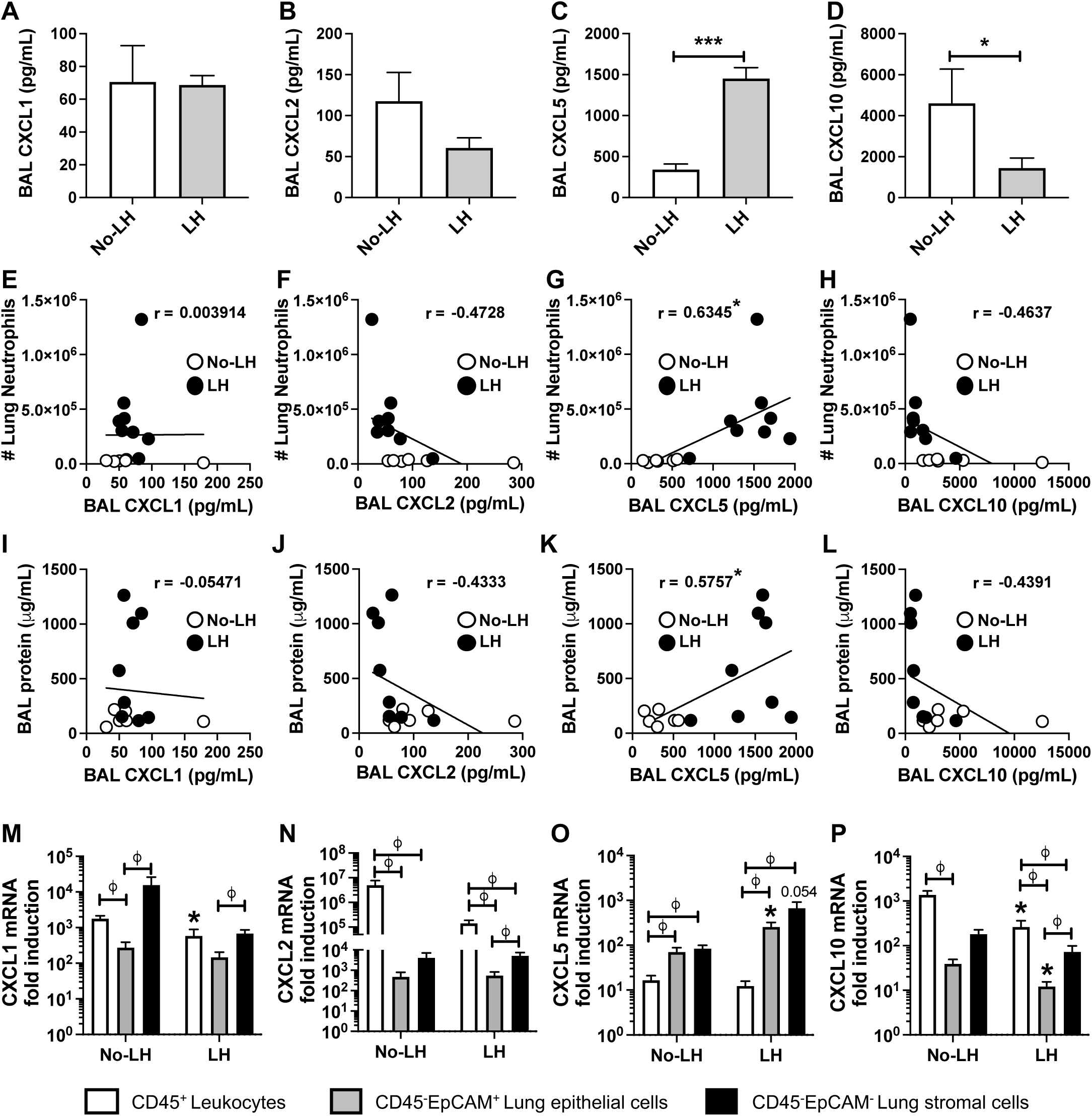
CXCL5 produced by lung epithelial and stromal cells correlates with severity of allergic airways neutrophilia. **A)** CXCL1, **B)** CXCL2, **C)** CXCL5 and **D)** CXCL10 levels in BALs of mice 24 hours post OVA challenge. *p* value: ***≤ 0.001 by Mann-Whitney test. **E-H)** Scatterplots correlating lung neutrophil numbers with **E)** BAL CXCL1, **F)** BAL CXCL2, **G)** BAL CXCL5 and **H)** BAL CXCL10 levels in mice 24 hours post OVA challenge. Spearman’s correlation coefficient (r) and statistical significance denoted. *p* value: *≤ 0.05. **I-L)** Scatterplots correlating lung damage (BAL protein) with **I)** BAL CXCL1, **J)** BAL CXCL2, **K)** BAL CXCL5 and **L)** BAL CXCL10 levels in mice 24 hours post OVA challenge. Spearman’s correlation coefficient (r) and statistical significance denoted. *p* value: *≤ 0.05. **M-P)** mRNA levels of **M)** CXCL1, **N)** CXCL2, **O)** CXCL5 and **P)** CXCL10 transcripts in FACS-sorted CD45^+^ leukocytes (*white*), CD45^-^EpCAM^+^ lung epithelial cells (*gray*), CD45^-^EpCAM^+^ structural cells (*black*) from mice 24 hours post OVA challenge expressed as fold change induction over naïve whole lungs. *p* value: ^Φ^≤ 0.05 by Kruskal-Wallis test between cell types but same treatment group, and *≤ 0.05 by Unpaired t test within a cell type but between treatment groups. All data have n≥5 mice, 2 experiments, mean ± SEM.

**Figure S6:**
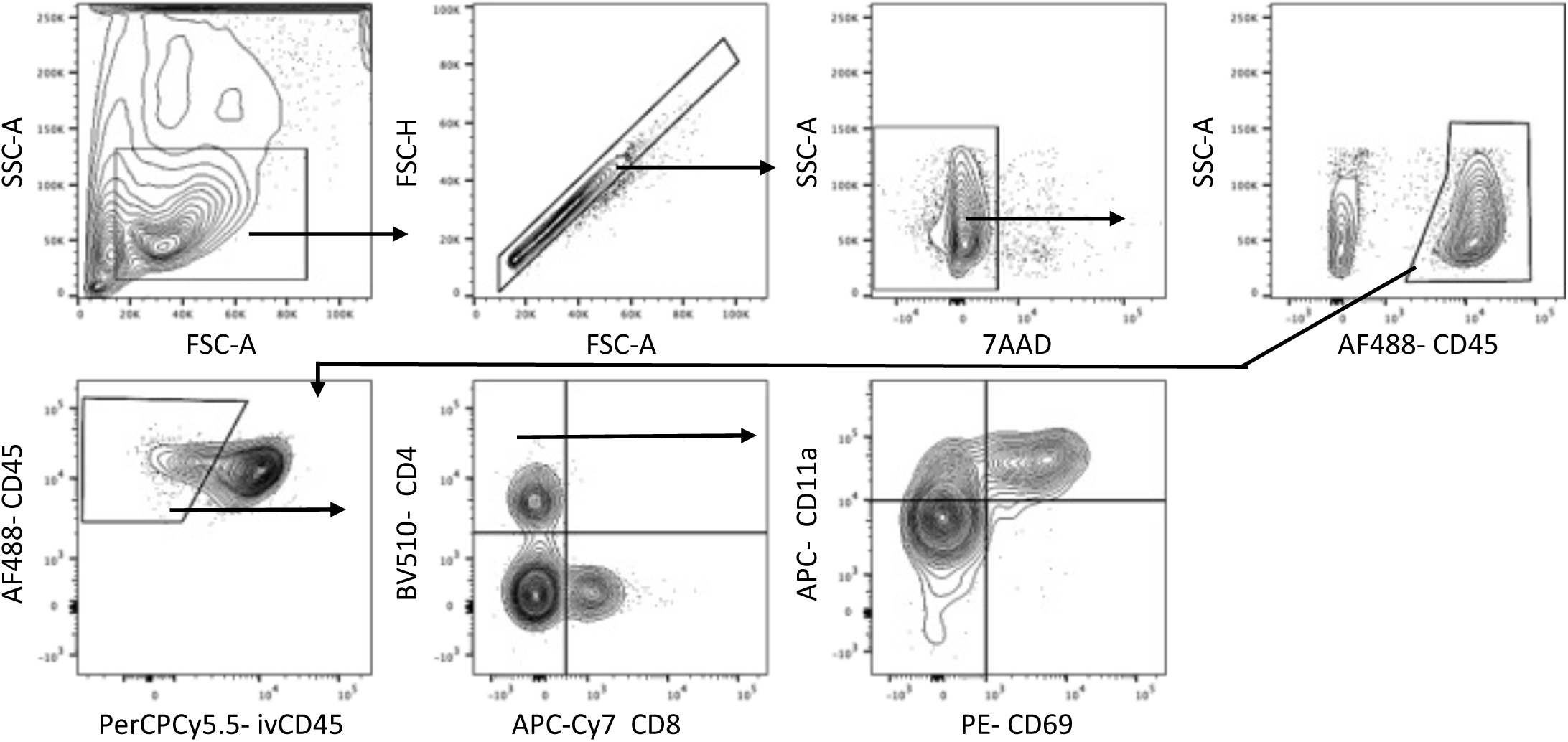
Gating strategy for identification of lung-resident CD4^+^ T_RM_ cells.

**Figure S7:**
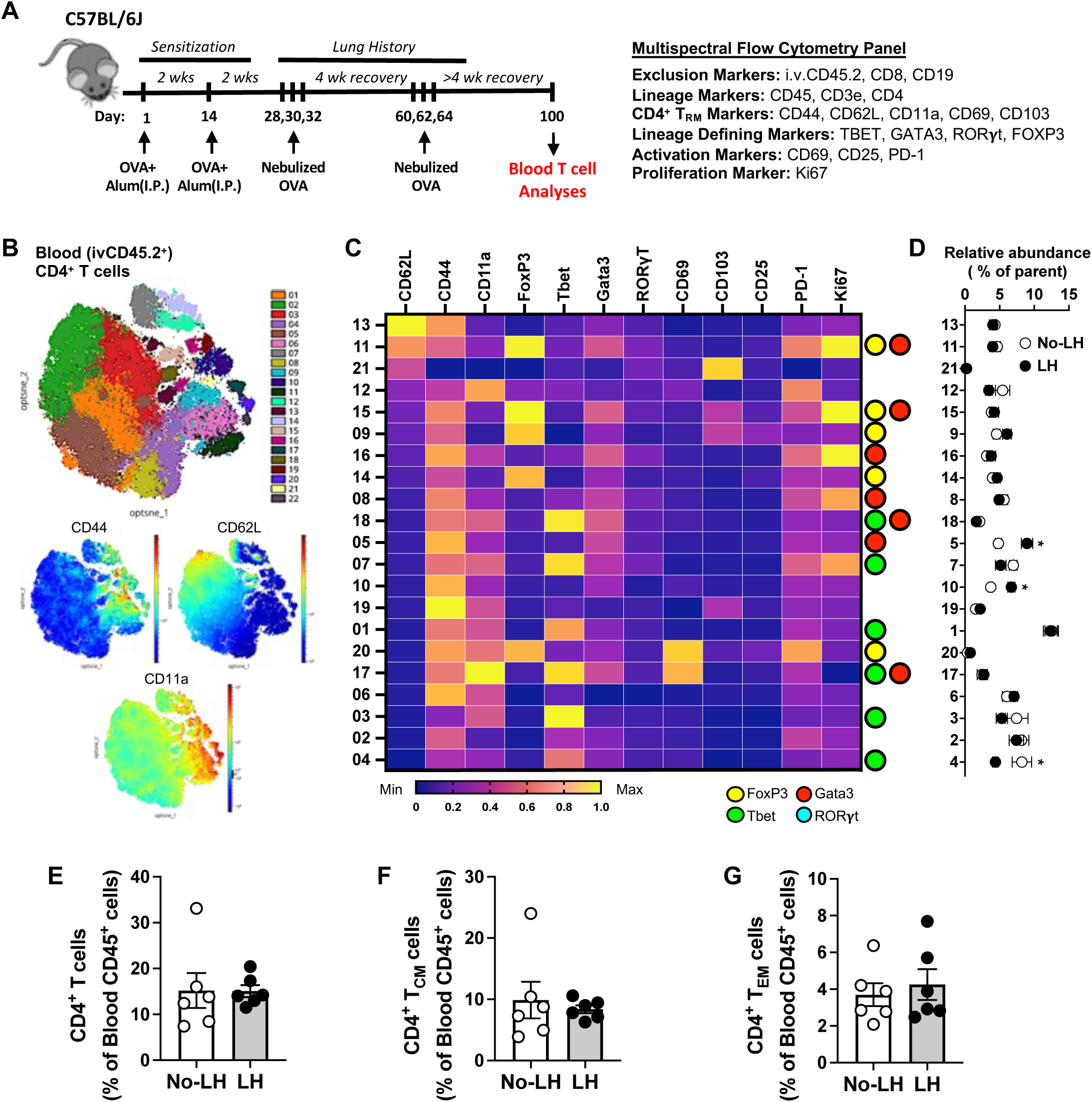
LH mice lack RORγt^+^ T_H_17 cells within circulation and share a comparable blood CD4^+^ T cell pool with No-LH mice. **A)** Schematic of experimental timeline and antibody panel used. **B)** Phenograph clustering overlaid on opt-SNE projection depicting blood (i.v.CD45.2^+^) CD4^+^ T cells concatenated from n=8 LH mouse lungs across 2 experiments on day 100. opt-SNE projection with heatmap visualization depicting CD44, CD62L and CD11a expression levels are shown. **C)** Heat map depicting normalized expression levels of distinct molecules on blood (ivCD45.2^+^) CD4^+^ T cell clusters. Lineage determining transcription factor (LDTF) status of each cluster is depicted on the *right*. **D)** Relative abundances of each blood (ivCD45.2^+^) CD4^+^ T cell cluster in LH mice (white) and No-LH mice(black). Two Way ANOVA with two stage step-up method of Benjamini, Krieger, and Yekutieli to correct for multiple comparisons. FDR *p* value: *≤0.05. Relative abundances of blood (ivCD45.2^+^) **E)** CD4^+^ T cells, **F)** CD62L^+^CD44^+^ CD4^+^ T_CM_ cells and **G)** CD62L^-^CD44^+^ CD4^+^ T_EM_ cells expressed as percentage fraction of blood (ivCD45.2^+^) CD45^+^ cells in LH mice (white) and No-LH mice (black). All data have n≥5 mice, 2 experiments, mean ± SEM.

**Figure S8:**
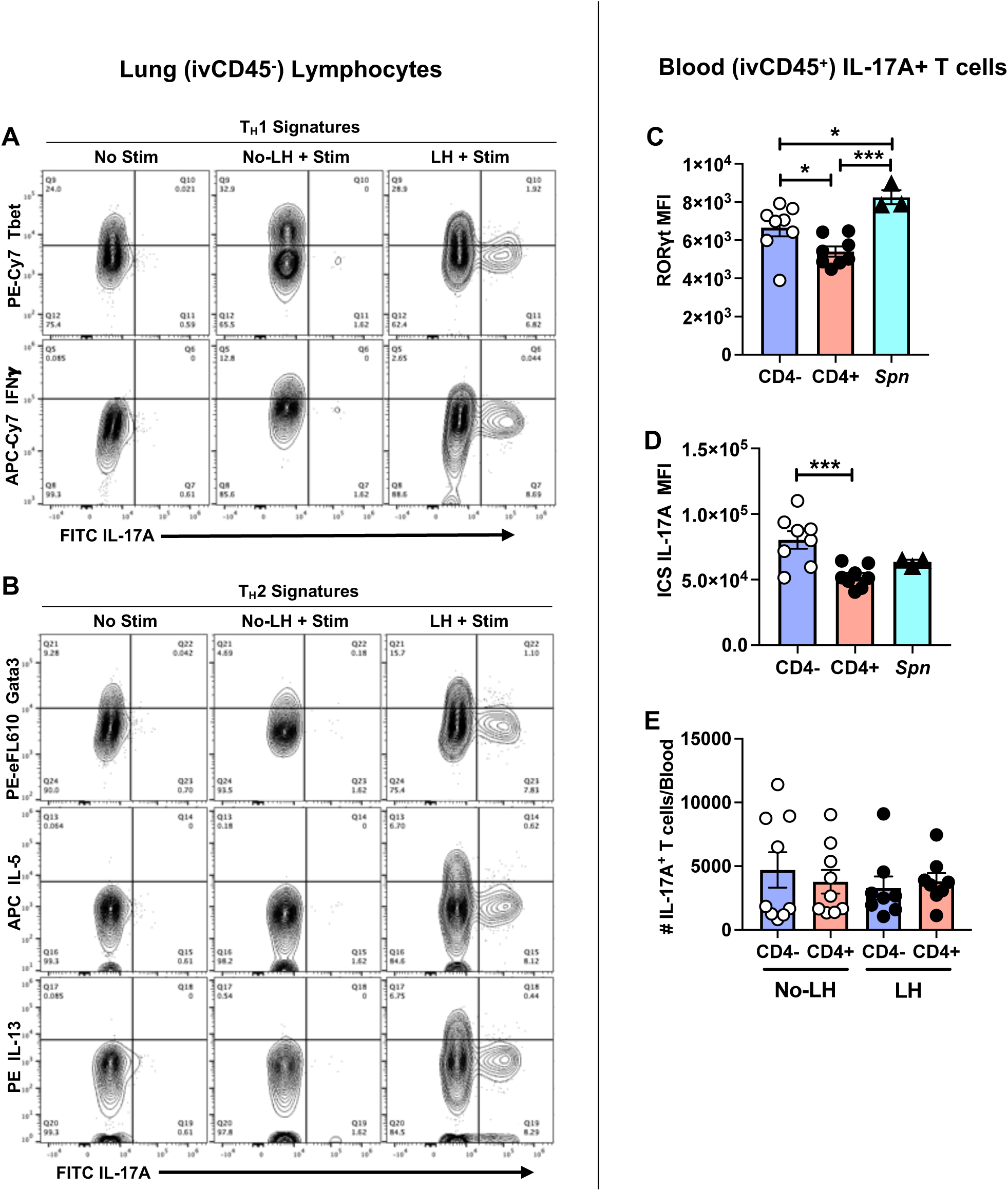
RORγt^negative/low^ T_H_17 T_RM_ cells are distinct from other traditional helper cell subsets and not enriched in blood. Representative contour plots depicting **A)** Tbet and IFN-γ, and **B)** Gata3, IL-5 and IL-13 expression patterns in RORγt^negative/low^ IL-17A^+^ producing lung (ivCD45.2^-^) CD4^+^ T cells in LH mice. Expression levels of **C)** RORγt and **D)** IL-17A in IL-17A^+^ blood (ivCD45.2^+^) CD4^-^ and CD4^+^ T cells in LH mice. One-way ANOVA with Fisher’s LSD Test. Expression patterns of *S. pneumoniae* induced (ivCD45.2^+^) CD4^+^ T cells are included as positive control for T_H_17 signatures. **J)** Absolute numbers of IL-17A^+^ blood (ivCD45.2^-^) CD4^-^ and CD4^+^ T cells in No-LH and LH mice. One-way ANOVA with Fisher’s LSD Test. All data have n≥3 mice, 2 experiments, mean ± SEM. *p* value: *≤0.05, **≤ 0.01, ***≤ 0.001.

**Figure S9:**
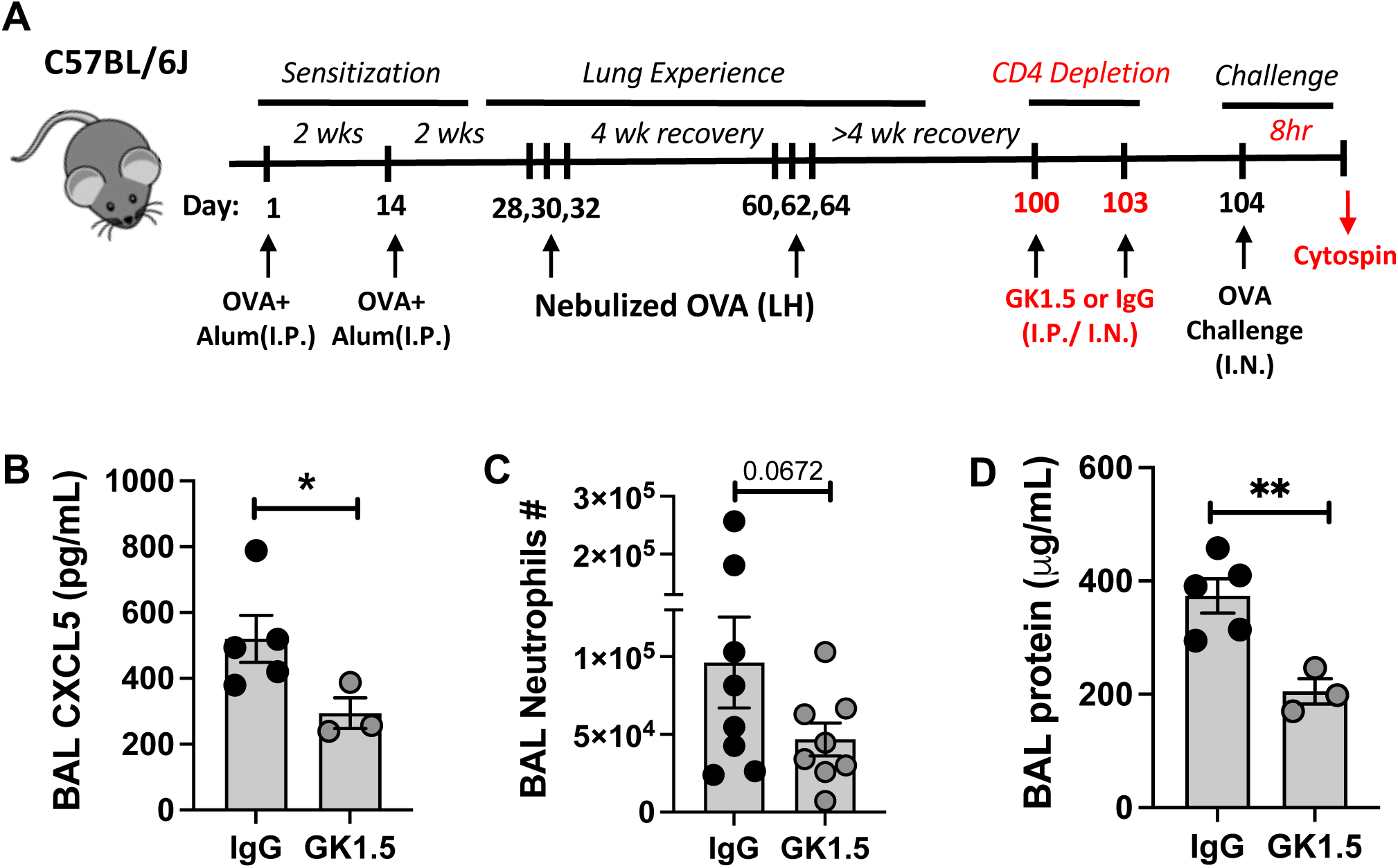
Depletion of CD4^+^ T_RM_ cells preempts allergic airway neutrophilia. **A)** Schematic of experimental timeline used. **B)** CXCL5 levels, **C)** neutrophils numbers, and **D)** edema in BAL of GK1.5 (or IgG) treated LH mice 8-hours post OVA challenge. Data except **S9B** and **S9D** has n≥5 mice, 2 experiments, mean ± SEM, One-tailed Unpaired t-test. *p* value: *≤ 0.05, **≤ 0.01.

**Figure S10:**
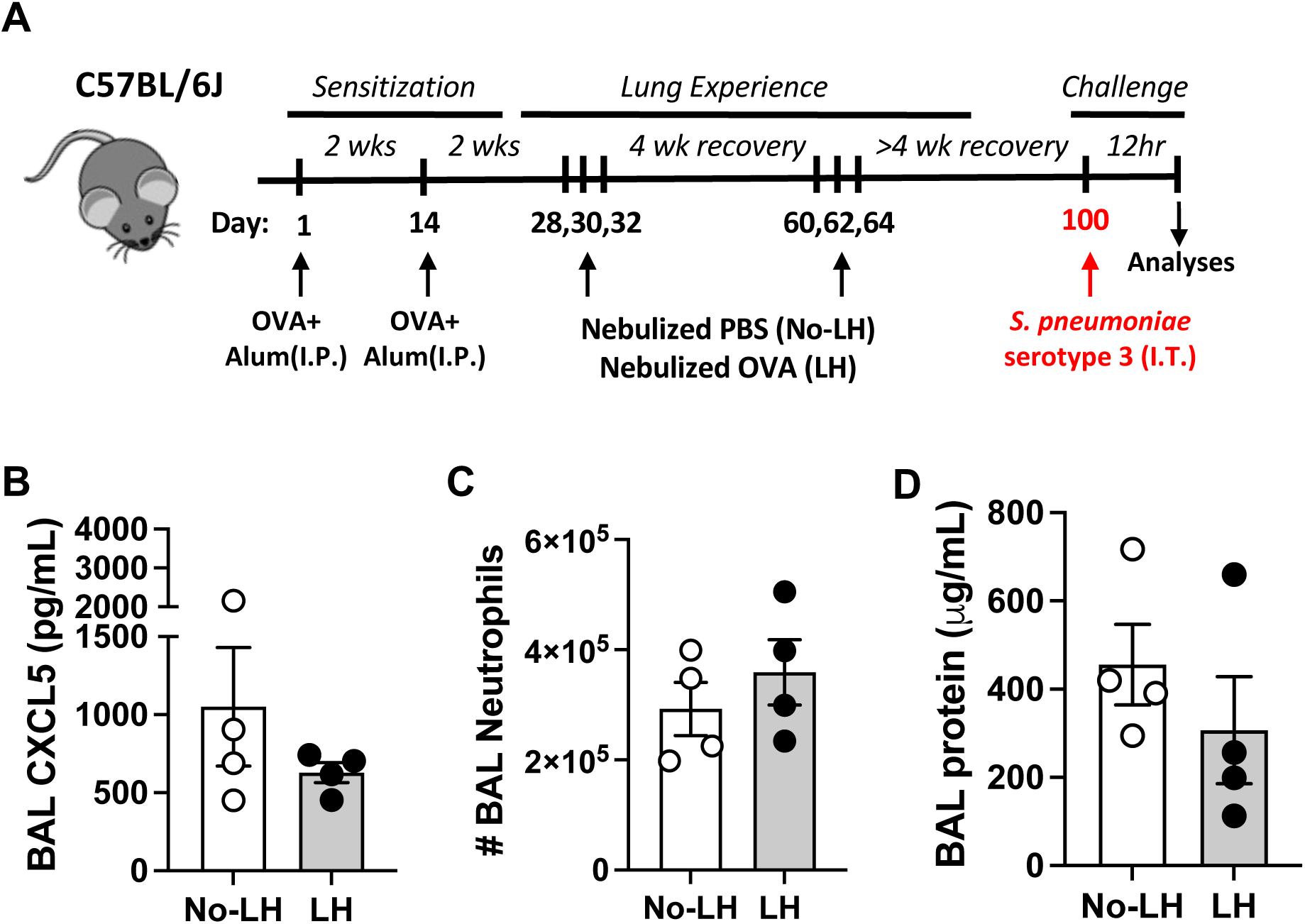
Allergic airway neutrophilia in LH mice is antigen specific and independent of trained immunity. **A)** Schematic of experimental timeline used. **B)** CXCL5 levels, **C)** neutrophils numbers, and **d)** edema in BAL of No-LH and LH mice 12-hours post *Streptococcus pneumoniae* serotype 3 challenge. *S. pneumoniae* was used as a model antigen irrelevant to the allergen to test role of trained immunity in the neutrophilic response. All data n=4 mice, 2 experiments, mean ± SEM.

**Figure S11:**
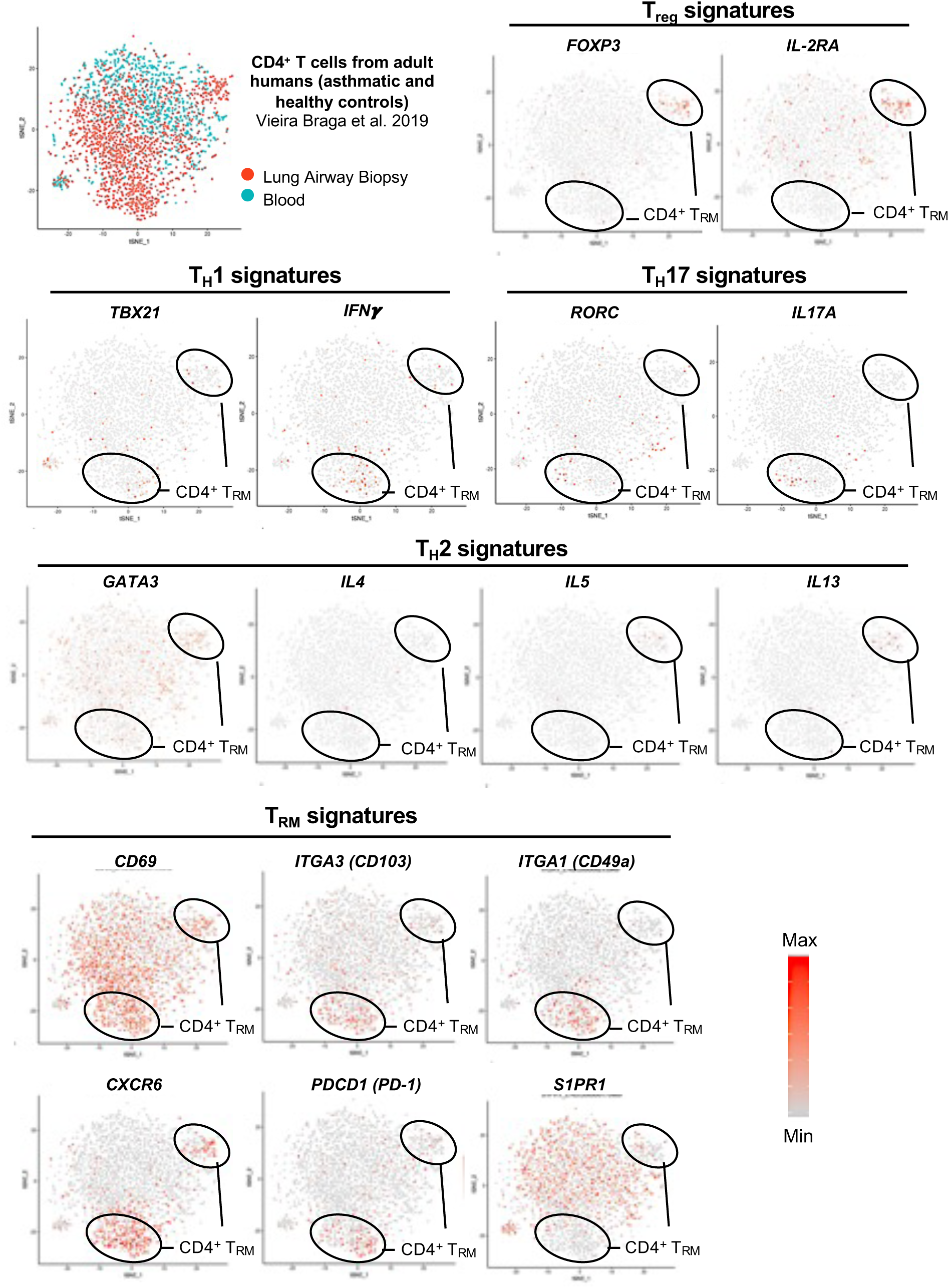
Airways of adult asthmatic and healthy human lungs harbor diverse clusters of CD4^+^ T_RM_ cells including RORγt^negative/low^ T_H_17 T_RM_ cells. t-SNE projection of scRNA-Seq data depicting expression patterns for diverse markers of interest as expressed by CD4^+^ T cell isolated from airway wall biopsies and peripheral blood of adult asthmatic and healthy humans and as displayed on the interactive web portal: https://asthma.cellgeni.sanger.ac.uk/ ^14^.

**Figures S12:**
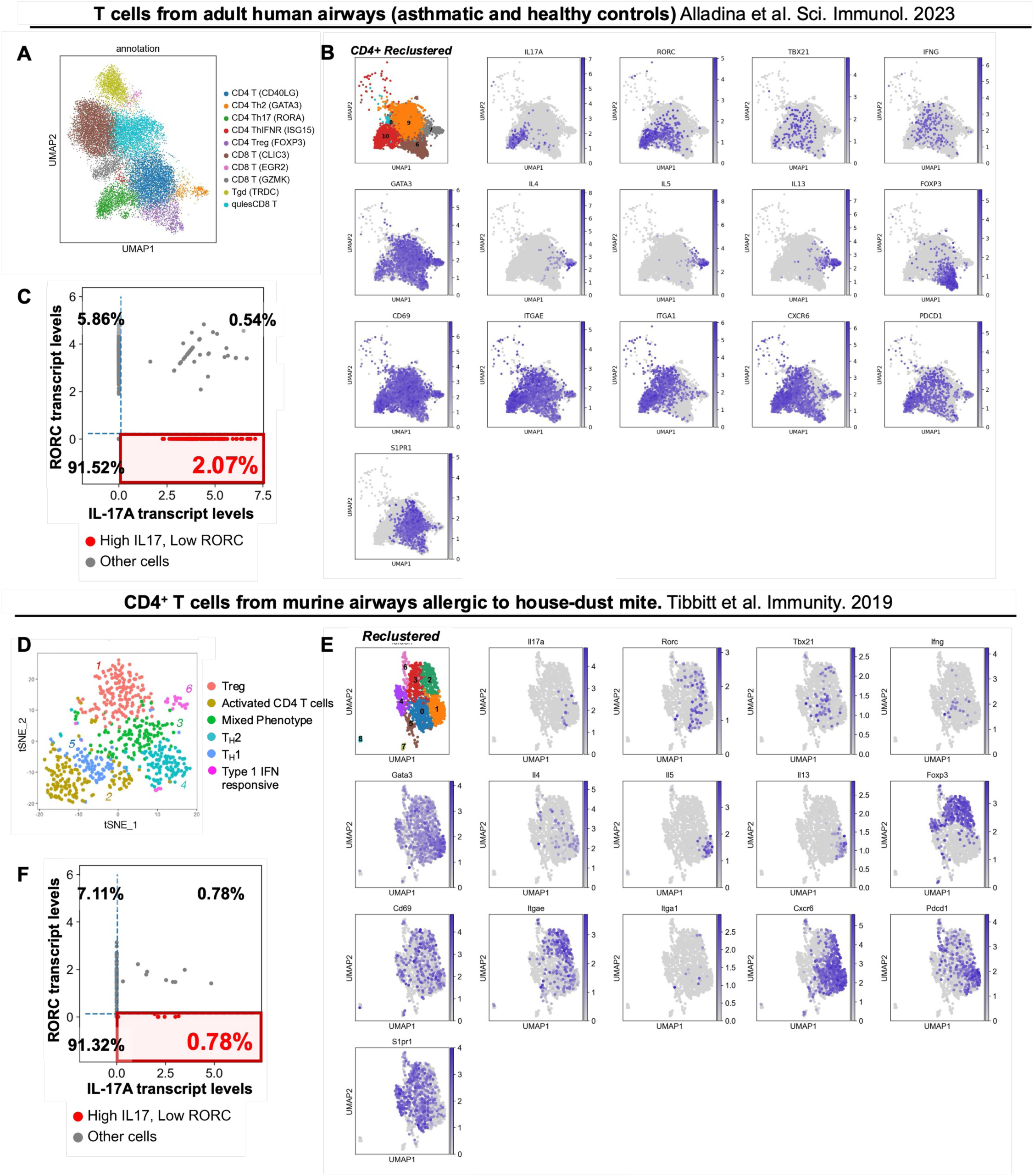
Airways of adult asthmatic and healthy humans (but not mice with allergic airways eosinophilia) are enriched for RORγt^negative/low^ T_H_17 T_RM_ cells. **A)** UMAP projection of scRNA-Seq data- and **B)** expression patterns for diverse markers of interest-as expressed by CD4^+^ T cell isolated from airway brushings of adult asthmatic 24 hours post segmental allergen challenge. **C)** Dot plot depiction of scRNA-Seq data showing frequencies of RORγt^negative^ T_H_17 T_RM_ cells in human airway brushings 24 hours post segmental allergen challenge. **Fig S12A-C** is obtained by reanalyses of published dataset^70^. **D)** UMAP projection of scRNA-Seq data- and **E)** expression patterns for diverse markers of interest-as expressed by CD4^+^ T cell isolated from airways (BAL) of mice exposed to HDM (house dust mites) induced allergic airways eoisnophilia. **F)** Dot plot depiction of scRNA-Seq data showing frequencies of RORγt^negative^ T_H_17 T_RM_ cells in mice with HDM-induced allergic airways eosinophilia. **Fig S12D-F** is obtained by reanalyses of published dataset^17^.

**Figure S13:**
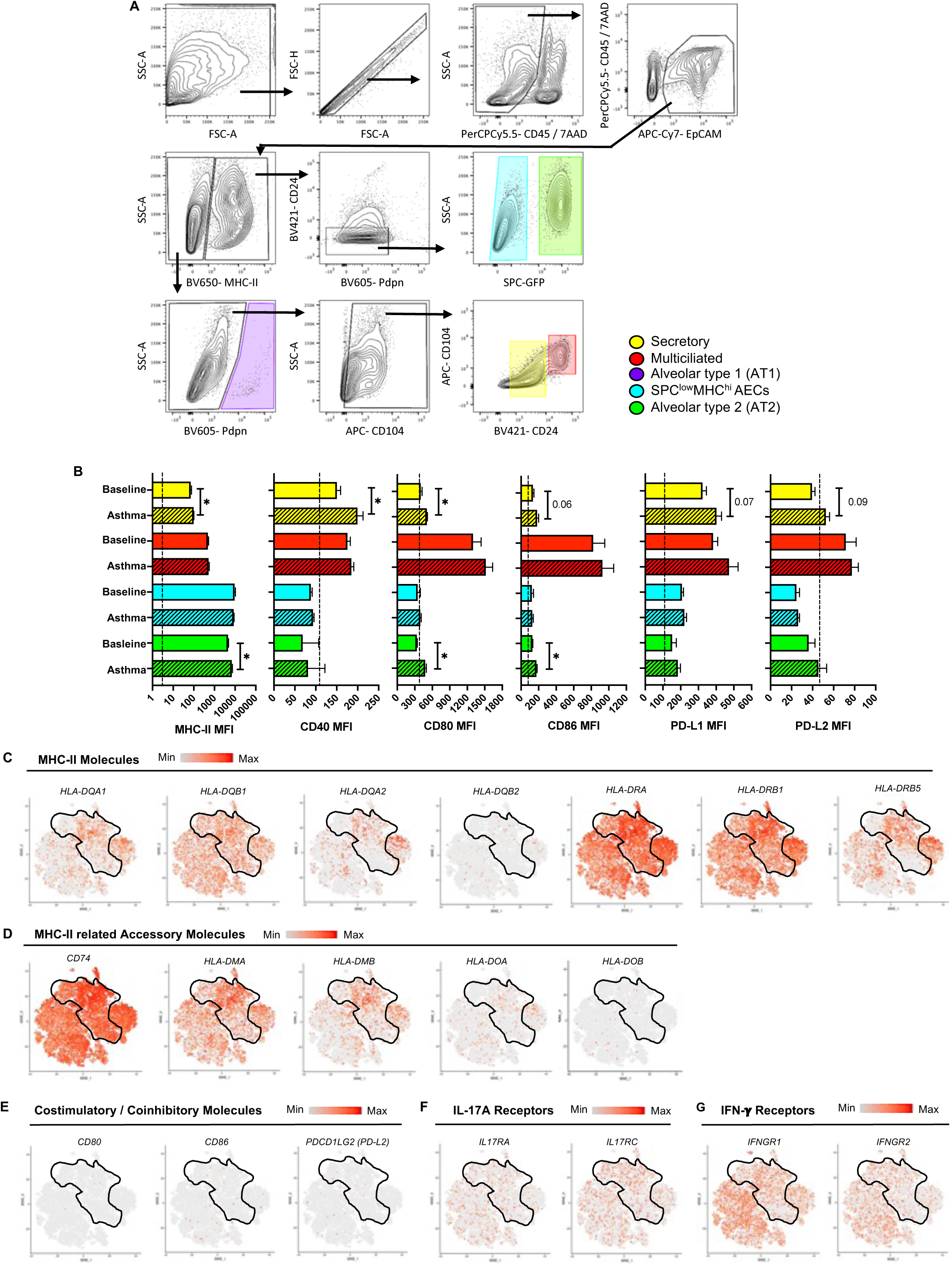
Airway Muc5ac^high^ secretory cells can communicate with CD4^+^ T_RM_ cells in lungs with extensive inhaled allergen history. **A)** Gating strategy for identification of distinct lung epithelial cell (LEC) subsets in SPC-GFP LH mice. **B)** Surface expression levels of specified APC-related molecules on distinct epithelial cells from LH mice at baseline (clear bars) and 24 hours post OVA challenge (slashed bars). Unpaired t test. Dotted lines indicate signal level on FMO controls. Mann-Whitney test. *p* value: *≤ 0.05. All data have n=4 mice, 2 experiments, mean ± SEM. **C-G**) t-SNE projection of scRNA-Seq data depicting expression patterns for designated markers of interest by different subsets of epithelial cells identified in airways of adult asthmatics and healthy humans as displayed on the interactive web portal: https://asthma.cellgeni.sanger.ac.uk/ ^14^.

**Figure S14:**
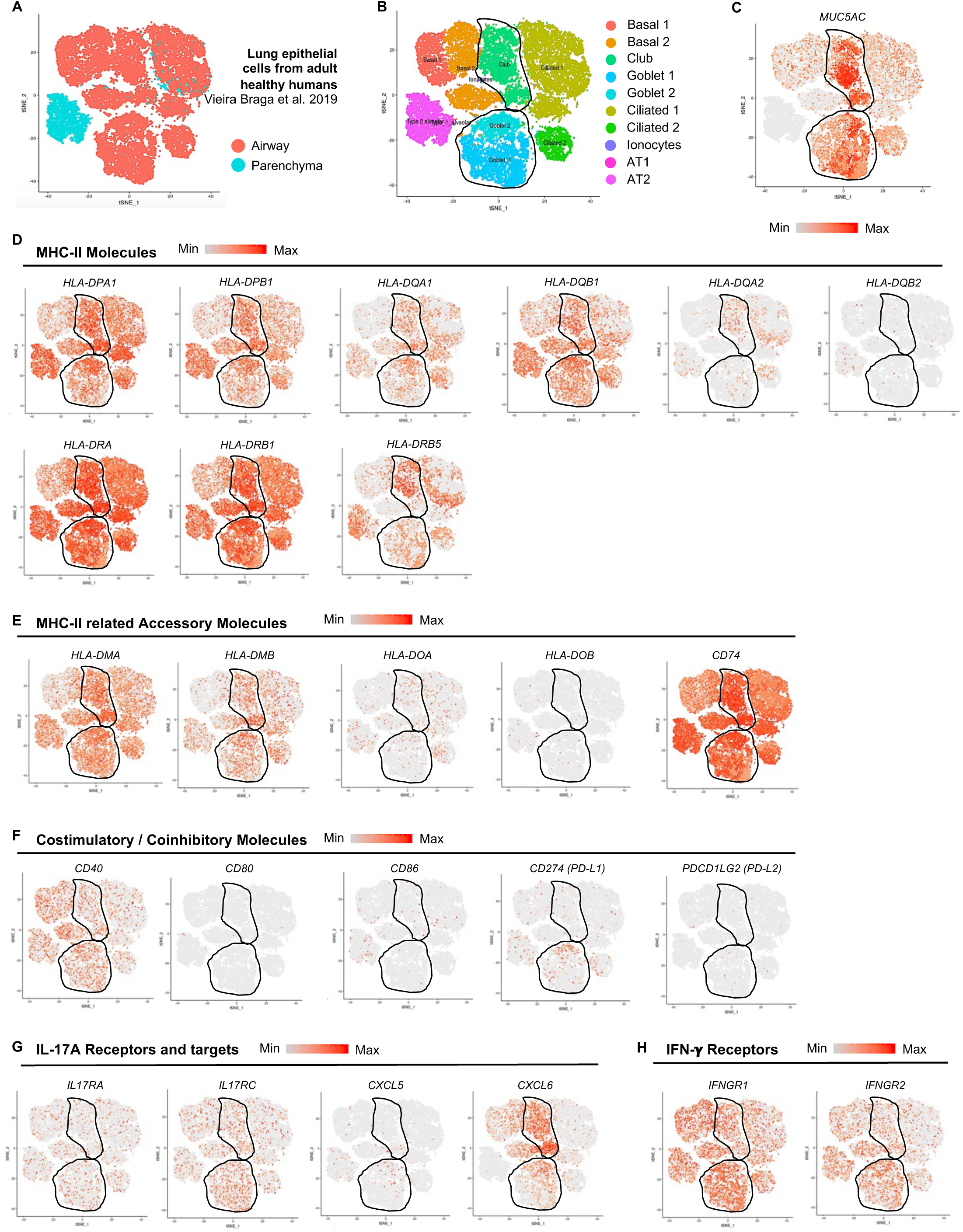
Muc5ac^high^ airway secretory cells are high expressors of CD4^+^ T cell and neutrophil facing molecules. t-SNE projection of scRNA-Seq data depicting expression patterns for designated markers of interest by different subsets of epithelial cells identified in airways and lung parenchyma of adult humans as displayed on the interactive web portal: https://asthma.cellgeni.sanger.ac.uk/ ^14^.

**Figure S15:**
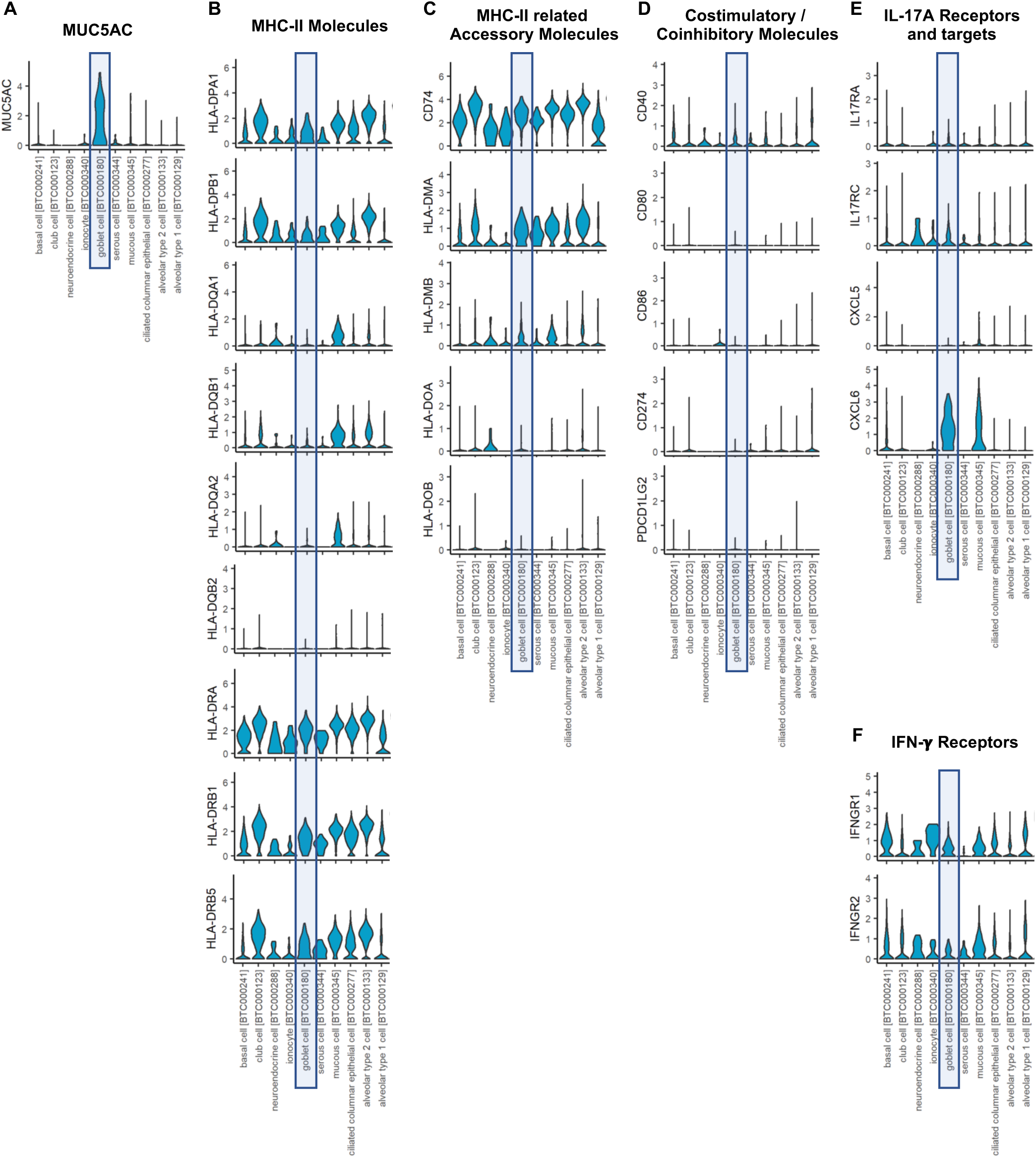
Muc5ac^high^ airway secretory cells may bridge CD4^+^ T_RM_ cells and neutrophils in lungs with extensive inhalation histories. t-SNE projection of scRNA-Seq data depicting expression patterns for designated markers of interest by different subsets of epithelial cells identified in lungs of adult humans ^73^.

**Figure S16:**
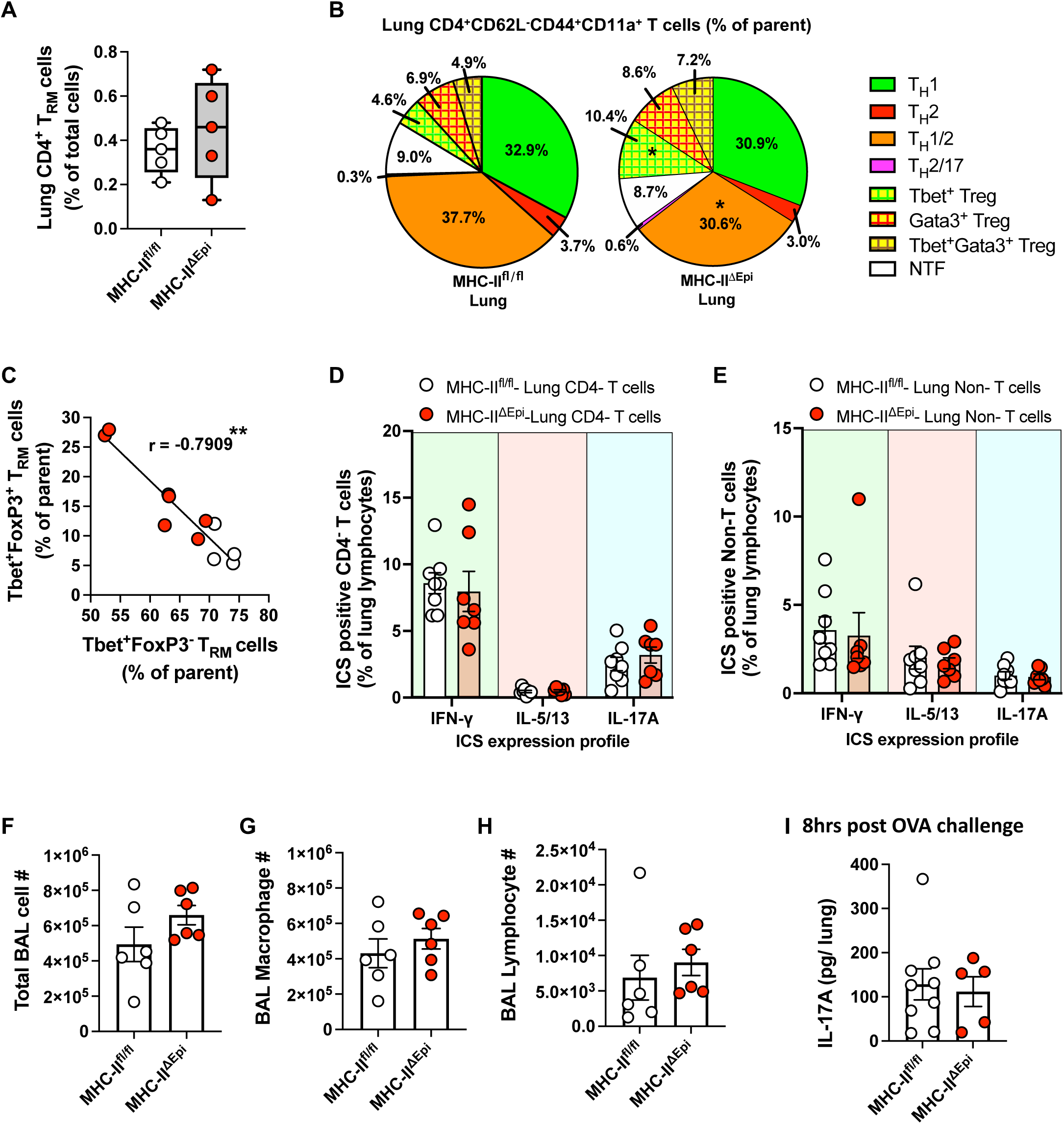
Characterization of LH mice lacking MHC-II specifically in lung epithelial cells. **A)** Abundance of lung CD4^+^ T_RM_ cell numbers at baseline. **B)** Pie charts for mean frequencies of distinct memory T_H_ cell lineages within lungs of MHC-II^fl/fl^ and MHC-II^ΔEpi^ LH mice at day 120, Two-way ANOVA with Fisher’s LSD test. p-value: *≤ 0.05. **C)** Scatterplot correlating fraction all Tbet^+^ FoxP3^-^ CD4^+^ T_RM_ cells with Tbet^+^ FoxP3^+^ T_RM_ cells. Spearman’s correlation coefficient (r) and statistical significance denoted. Positivity for a tested LDTF was determined using cutoffs identified from clusters negative for that LDTF. **D, E)** Intracellular cytokine staining (ICS) profile of **D)** lung (i.v.CD45.2^−^) CD4^-^ T cells and **E)** lung (i.v.CD45.2^−^) CD3^-^ non-T cells isolated from lungs on day 120 and stimulated with PMA/Ionomycin *ex vivo*. Two-way ANOVA with Fisher’s LSD Test. **F)** BAL cellularity-, **G)** BAL macrophage-, and **H)** BAL lymphocyte-numbers at 24 hours post OVA challenge. **I)** Levels of whole lung IL-17A in mice 8 hours post OVA challenge. All data have n≥4 mice, 2-3 experiments, mean ± SEM.

**Figure S17:**
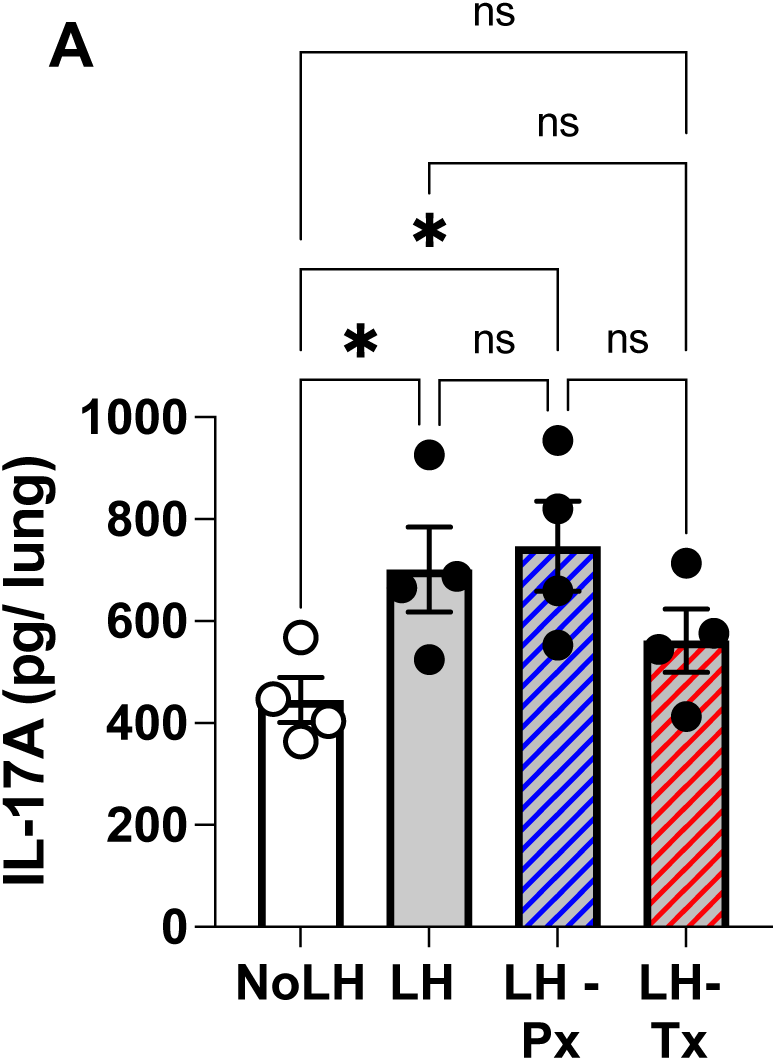
Exogenous supplementation of IFN-γ does not affects IL-17A levels in LH mice. A) Levels of whole lung IL-17A in mice 24 post OVA challenge with specified treatment modalities. Ordinary One-Way ANOVA with Fisher’s LSD Test. All data have n=4 mice, 2 experiments, mean ± SEM. p-value: *≤ 0.05.

**Figure S18:**
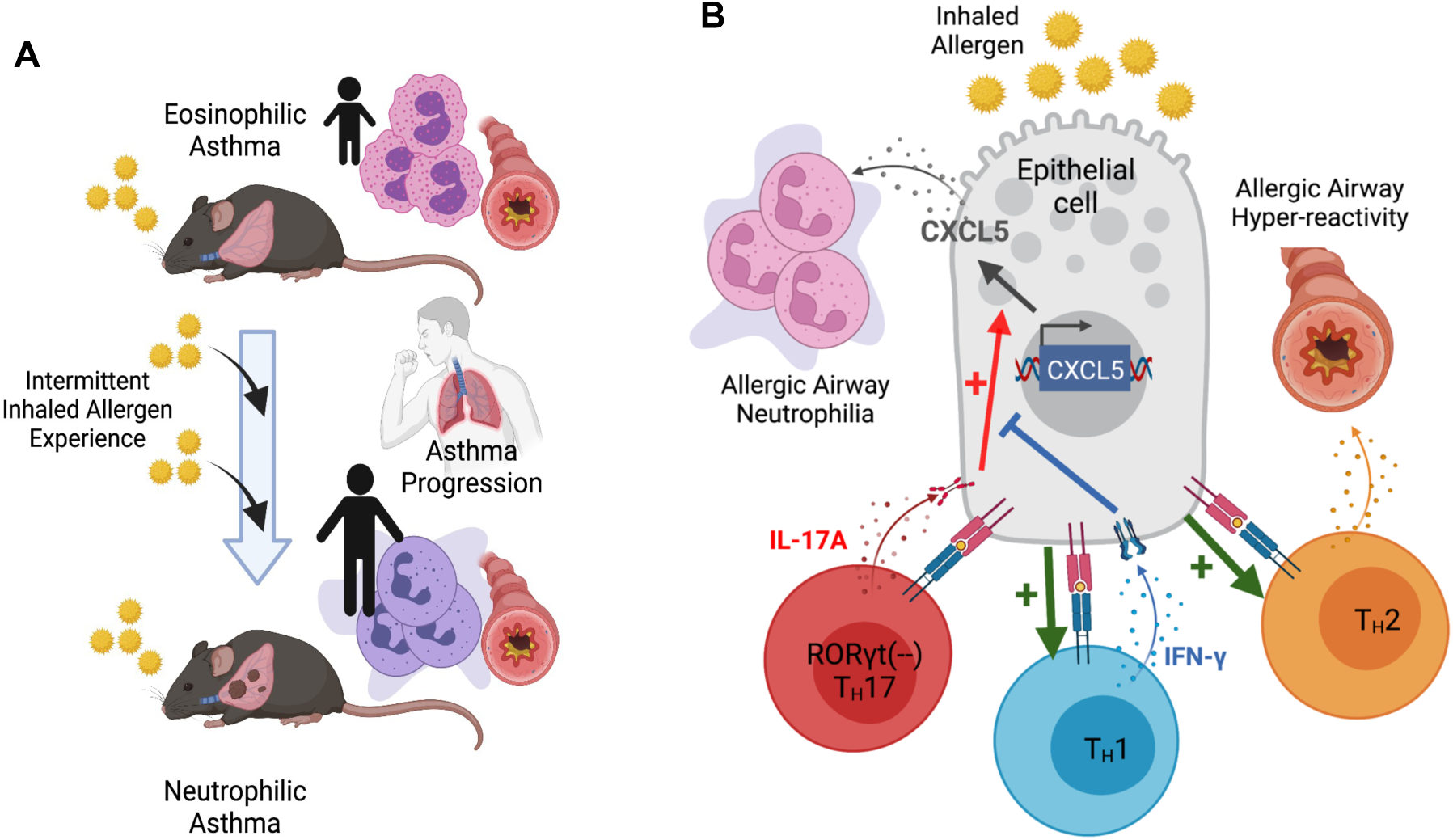
Proposed model. **A)** Intermittent exposures of sensitized mice to inhaled allergens over extended duration induces progression of eosinophilic allergic airways disease to neutrophilic allergic airways disease that recapitulates hallmark features of destructive neutrophilic asthma. **F)** Lungs of mice with age-appropriate inhaled allergen exposure history harbor diverse clusters of tissue-resident CD4^+^ T_RM_ cells which includes T_H_1, T_H_2, and unconventional RORγt^neg/low^ T_H_17 T_RM_ cells. Antigenic reactivation of the latter leads to secretion of IL-17A, inducing expression of epithelial CXCL5 which elicits peribronchial neutrophilic infiltration in association with more severe disease. Antigen presentation by epithelial MHC-II bolsters T_H_2 T_RM_ cells augmenting airway hyper-reactivity, and T_H_1 T_RM_ cells; the latter via its effector cytokine IFN-γ curbs CXCL5 and airway neutrophilia driven by IL-17A. Thus, lung epithelial cells instruct CD4^+^ T_RM_ cell fates and activities to control the type and extent of airway allergic responses.

## Notes

### Competing Interest Statement

The authors have declared no competing interest.

